# Cerebrospinal Fluid Flow Extends to Peripheral Nerves

**DOI:** 10.1101/2023.11.20.567884

**Authors:** Alexander P. Ligocki, Augustine V. Vinson, Anthony T. Yachnis, William A. Dunn, Douglas E. Smith, Elizabeth A. Scott, Jimena V. Alvarez-Castanon, Daniel E. Baez Montalvo, Olivia G. Frisone, Gary A.J. Brown, Joel E. Pessa, Edward W. Scott

## Abstract

Cerebrospinal fluid (CSF) is an aqueous solution responsible for nutrient delivery and waste removal for the central nervous system (CNS). The three-layer meningeal coverings of the CNS support CSF flow. Peripheral nerves have an analogous three-layer covering consisting of the epineurium, perineurium, and endoneurium. Peripheral axons, located in the inner endoneurium, are bathed in “endoneurial fluid” similar to CSF but of undefined origin. CSF flow in the peripheral nervous system has not been demonstrated. Here we show CSF flow extends beyond the CNS to peripheral nerves in a contiguous flowing system. Utilizing gold nanoparticles, we identified that CSF is continuous with the endoneurial fluid and reveal the endoneurial space as the likely site of CSF flow in the periphery. Nanogold distribution along entire peripheral nerves and within their axoplasm suggests CSF plays a role in nutrient delivery and waste clearance, fundamental aspects of peripheral nerve health and disease.

**One Sentence Summary:** Cerebrospinal fluid unites the nervous system by extending beyond the central nervous system into peripheral nerves.

## INTRODUCTION

Cerebrospinal fluid (CSF) is an ultrafiltrate of the plasma produced by the Choroid plexus found in the ventricles of the brain, that bathes the tissues of the central nervous system (CNS). CSF is continuously produced (approximately 0.5 L per day in humans), providing physical cushioning and suppling nutrients, while points of efflux from the system are vital for removing waste (1–10). In essence the CSF flow system is critical for maintaining homeostasis of nervous tissues. In addition to the critical role of CSF in maintaining homeostasis in the central nervous system, the possible passage of CSF or its’ constituents to the peripheral nervous system has been suggested(10–14).

After production by the ependymal cells lining the choroid plexus and the ventricles of the brain, CSF travels via low pressure pulsatile flow within the central nervous system. Outside the ventricles, the CSF is physically contained within the subarachnoid space (SAS) of the brain and spinal cord, a space created between the pia mater and the arachnoid mater (4). The dura is the dense, outermost layer of meninges attached to the skull containing venous sinuses and lymphatics(2, 6, 15, 16). Peripheral nerves have an analogous three-layer connective tissue covering that is continuous with CNS meninges. The epineurium (analogous to the dura) bundles individual nerve fascicles that are enclosed by the perineurium (analogous to the arachnoid). This enclosed region contains the endoneurium, a loose connective tissue space that surrounds the bundled axons, creating a privileged environment needed for optimal nerve functioning (17, 18). The endoneurium surrounding the Schwann cells that covers peripheral axons contains fluid similar in composition to CSF, yet the origins of this fluid are debated(17, 18). Previous vital dye studies showed dye collecting in a cuff around the roots of peripheral nerves where they emerge from the CNS following infusion into the subarachnoid spaces, but to date no conclusive evidence of CSF flow throughout the peripheral nervous system (PNS) has been demonstrated(2, 10–14, 17).

In this study we infused 1.9nm gold particles (nanogold) into a lateral cerebral ventricle, the major site of CSF production, to demonstrate continuous CSF flow from the sites of production in the CNS to peripheral nerves in live mice. Peak distribution of nanogold delivered as a single intracerebral ventricular (ICV) bolus was size, time, and concentration dependent. CNS structures were labeled within minutes, while delivery to the distal sciatic nerve peaked between 4-6 hours (hr) post infusion. Electron microscopy confirmed nanogold delivery from the cerebral ventricle to the axoplasm of distal peripheral nerves. Our results support a contiguous and continuous CSF flow system from the CNS to the distal ends of peripheral nerves that is likely integral in sustaining peripheral nerves through delivery of nutrients and removal of waste.

## RESULTS

### Ventricular Infusion of Gold Nanoparticles Demonstrates CSF Flow to Peripheral Nerves

The 1.9 nm nanogold was selected as a tracer due to its size being similar to many of the nutrients carried by CSF(19). Furthermore, this nanogold can be detected by multiple imaging modalities which utilize enhanced staining methods to increase sensitivity of detection for visualization via light and electron microscopy. A single bolus of 1.9 nm nanogold was infused via a fine cannula within the lateral ventricle of a sedated mouse at a rate demonstrated to maintain physiologic distribution within the CNS (Fig. 1A-M, Fig.S1, S2)(5, 7, 20, 21).

**Figure 1.**
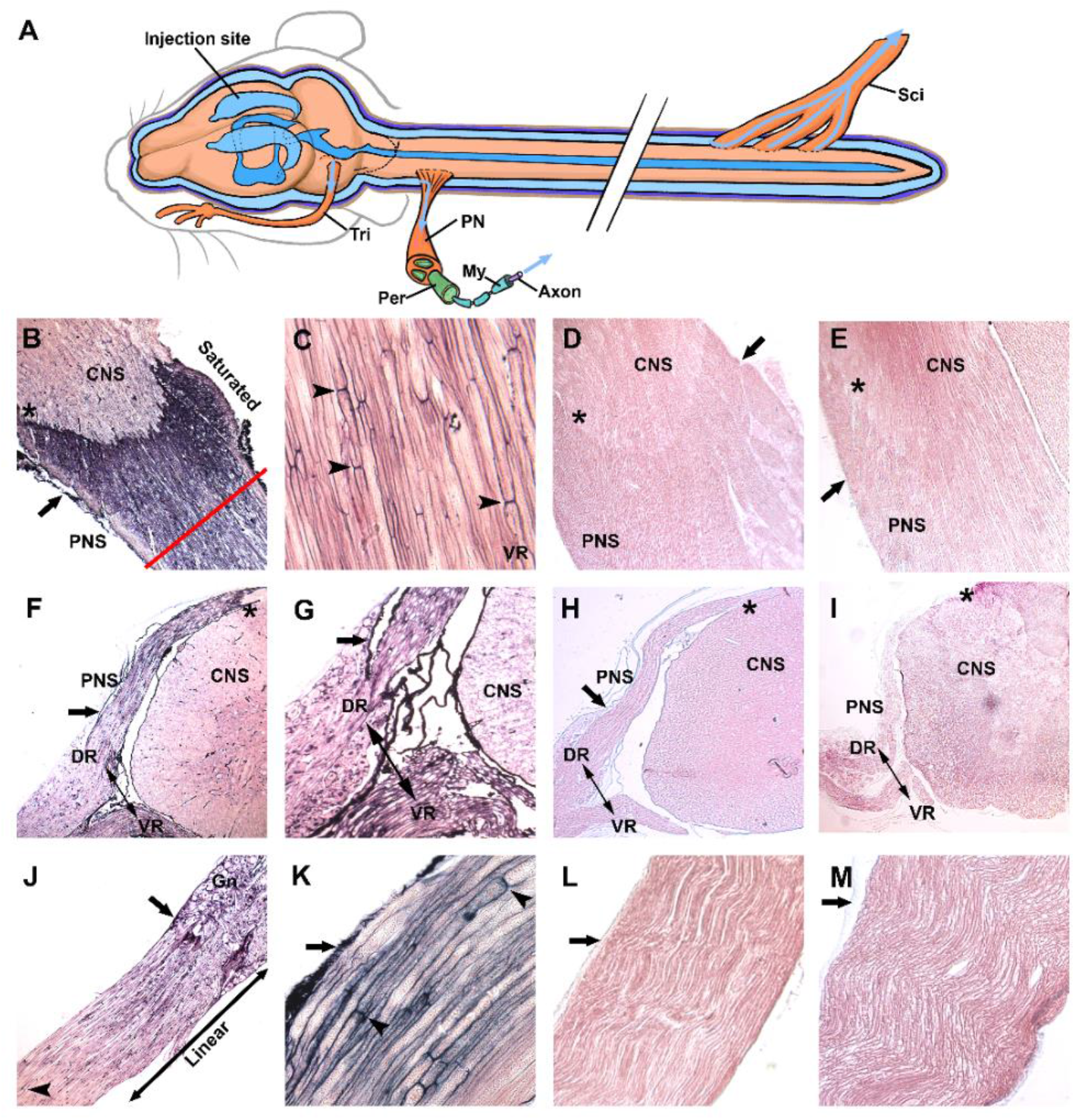
CSF infused nanoprobe travels to peripheral nerves. Peripheral nerves 4 - 6hr after nanoprobe ICV-infusion (B,C,F,G,J,K), IV infusion (E, I, M) and PBS controls (D,H,L). Trigeminal nerve (B-E), spinal cord (F-I) and sciatic nerve sections (J-M) with black GoldEnhance staining. CNS-PNS transition marked by “*” in all panels. (A) Schematic of CSF reservoirs; Peripheral nerve (PN), Axon (Ax), Myelin (My), Perineurium (Peri), Endoneurium (Endo). (B,C) Trigeminal nerve 4hr post infusion. (B) Staining shows CNS-PNS transition. Perineurium demarcated by closed arrow in all panels; arrowheads mark Nodes of Ranvier. (C) Staining is present in the endoneurium of trigeminal nerves, and Nodes of Ranvier (arrowhead). (D) PBS injected control. (E) Trigeminal nerve from IV animal 6hr after nanoprobe infusion. (F,G) Dorsal and ventral spinal nerve roots (double sided arrow) (NR) from nanoprobe-infused animals. Closed arrow marks subarachnoid angle of dorsal spinal nerve root. (G) Subarachnoid angle. (H) Spinal cord PBS-control. (I) Spinal cord cross section IV-injected animal 6hr post infusion. (J) Sciatic nerve taken at the ganglion 4hr after nanoprobe injection (J) and high magnification 6 hr after nanoprobe injection (K). Gold staining revealing Nodes of Ranvier. (L) PBS-control sciatic nerve. (M) 6hr. Sciatic nerve IV control. Minimum N=4.

To ensure our experimental design and probe faithfully trace CSF flow, we assessed whether our probe system recapitulated established CSF flow patterns in the CNS. To do this, we analyzed CNS tissues at immediately following 1.9 nm probe infusion (0 hr), a time when ICV injected tracers flow patterns has been well documented (Fig. S2) (1, 11, 12, 22). We examined the lateral ventricle infusion site immediately after the infusion was complete (Fig. S2A). Gold labeling was seen at the apical surface of the ependymal cells lining the ventricle and of the choroid plexus as well as along the Foramen of Monro that connects the lateral and third ventricle (Fig. S2A, B). During infusion, nanogold traveled to the dorsal third ventricle with robust labeling of the ependymal cells of the choroid plexus (Fig. S2B). Molecules injected into the CSF spaces are known to flow between meningeal layers in the subarachnoid space of the CNS(4, 8, 10). We saw significant labeling of the meninges of the brain and the optic nerve, the only cranial nerve considered as part of the CNS (Fig. S2C). Nanogold traversed the cerebral aqueduct (Fig. S2D) to the fourth ventricle then to the central canal of the spinal cord (Fig. S2E). These data demonstrate that nanogold follows previously established CSF flow patterns within the CNS(4, 8–10).

CSF flow was assessed by nanogold distribution in peripheral nerves, namely the trigeminal, spinal, and sciatic nerves (Methods, Experimental Details, and Supplementary Figures available online). We used a gold enhancement process (GoldEnhanceTM Nanoprobes, Yaphank,NY) to generate black gold aggregates visible by light and electron microscopy to assess CSF flow in the CNS and PNS (Fig 1). Sections of peripheral nerves harvested 6 hr after nanogold ICV infusion show nanogold distribution from the CSF of the central nervous system to peripheral nerves (Fig. 1B, C, F, G, J, K). Control tissues from PBS-ICV animals (Fig. 1D, H, L) show only background staining after gold enhancement (Fig. S3). Substantial deposition of nanogold was seen in the perineurium of peripheral nerves (Fig. 1B, F, J, K arrows), and deposition was observed within the endoneurium, resulting in distinctive labeling of the Nodes of Ranvier compared to control tissues (Fig. 1C, J, K arrowheads). To control for vascular influence on nervous system staining, we performed tail vein/intravenous (IV) injections with the same amount of nanogold used for ventricular injections. We then harvested the corresponding tissue at 6 hr, the same timepoints shown for PNS staining analysis (Fig. 1E, I, M). No nanogold staining above background was seen in peripheral nerves in IV nanogold-injected animals (Figure 1E, I, M). Low magnification image of trigeminal nerve (Fig. 1B) and nerve roots emerging from the spinal cord (Fig. 1F, G) showed significant labeling of the peripheral nerve compared to minimal nanogold deposition within the CNS parenchyma (Fig. S3). Differential staining intensity within the peripheral nerve root compared to the brain (Fig. 1B asterisk) spinal cord (Fig. 1F asterisk) in nanogold-infused tissues clearly marks the transition between central and peripheral nervous systems. Higher magnification of the cervical nerve roots revealed a lack of gold deposition at the subarachnoid angle; rather probe deposition was seen accumulating at the transition from CNS to PNS before flowing into the endoneurium surrounding individual axons (Fig. 1C, G, J, K). Sciatic nerve staining showed the highest concentration of gold in the perineurium (Fig. 1J, K), with clear labeling of endoneurium and the Nodes of Ranvier observed at 6 hr post infusion versus clear controls (Fig. 1D, H, L, E, I, M). Quantification of enhanced gold staining was determined by black pixel intensity using the Aperio ImageScope software, allowing identification of saturated (Fig. 1B red line) from linear (Figure 1J marked linear) and background areas of staining set by PBS control tissues (Fig.1D, H, L) confirmed higher probe deposition in the PNS compared to the CNS in all tissues (Fig. S3).

### Differential probe deposition patterns reveal size-dependence of CSF solute transport

Previous studies that define the meningeal lymphatics within the dura used 15nm fluorescent Q-dots and the glymphatic studies used a variety of tracers ranging from 2-3 nm to > 32 nm, however these studies did not assess transit to peripheral nerves(5, 6). Early studies on CSF flow used large particles such as India Ink, to evaluate flow into peripheral nerves (11, 12). To assess the effect of particle size on nanogold transport from CNS to PNS we infused 1.9 nm and 15nm nanogold into the lateral cerebral ventricle of mice as described above and analyzed probe deposition patterns in tissues 0hr and 4hr after infusion without gold enhanced staining (Fig. 2). Note, 1.9nm nanogold solution is black while 15nm nanogold solution is red via light microscopy without enhancement. Amplification with gold enhanced staining results in black deposits via light microscopy for both sizes of nanogold tracer. At 0hr post infusion overall levels of staining within the brain and SAS of transected spinal cord were similar for both nanogold sizes (Fig. 2A, C, E, G, I, K) versus PBS infused controls (Fig. S4). However, by 4hr post infusion, the amount of 15nm nanogold was reduced compared to 0hr post infusion, demonstrating clearance from the CNS (Fig. 2 C vs. D, G vs. H, K vs. L). In contrast, infused 1.9nm nanogold was retained within the whole brain with visible darkening of the parenchyma by 4hr post infusion (Fig. 2A vs. B). 1.9nm nanogold was also retained better within the spinal cord 4hrs post infusion (Fig. 2E vs. F, I vs. J). Both 0hr and 4hr infused 1.9nm axial view spinal cords show an almost uniform staining, continuing into peripheral nerves (Fig. 2I, J), whereas 0hr infused 15nm nanogold spinal cord axial views show accumulation of tracer around the emerging peripheral nerve roots (Fig. 2K). By 4hr post infusion spinal cord parenchyma has mostly cleared of 15nm nanogold with areas of tracer accumulation at the roots of emerging peripheral nerves remaining (Fig. 2L). This phenomenon was described in early CSF tracer studies and has been called “cuffing”(11, 12). Analysis of the emergent trigeminal nerves shows evidence of 1.9nm nanogold darkening of the nerve by 4hr post infusion (Fig. 2M vs. N). 15nm nanogold shows trigeminal nerve cuffing at both 0hr and 4hr post infusion (Fig. 2O, P).

**Figure 2.**
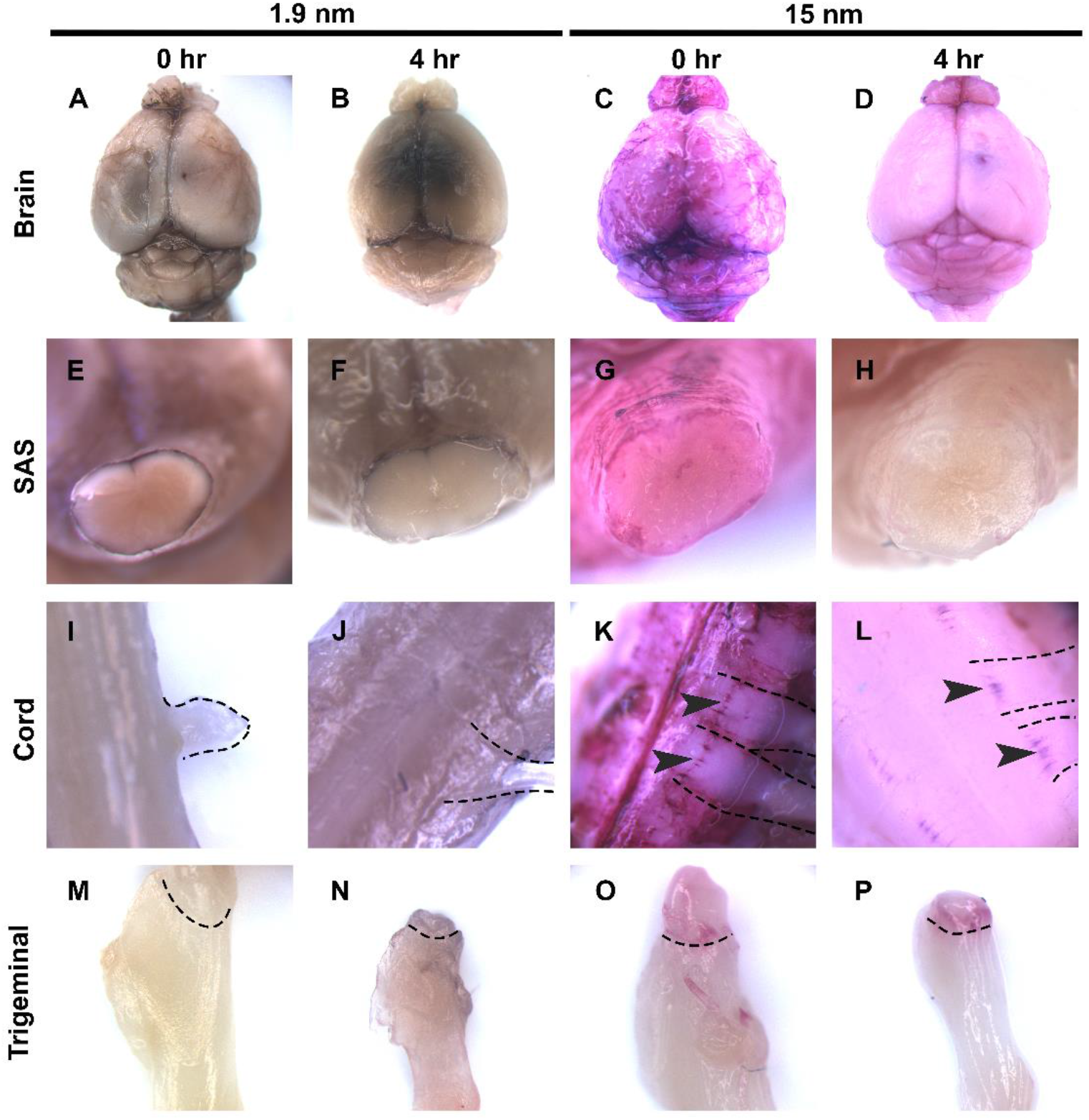
Larger Nanogold forms cuffing at CNS-PNS transitions. 1.9 or 15nm-gold tracer was infused into the lateral cerebral ventricle of fully anesthetized mice. CNS and PNS tissues were harvested 0hr or 4hr after ICV tracer infusion and imaged without gold enhancement. Whole brains were imaged with 1.9nm-gold (A) 0hr, (B) 4hr post infusion. Brains infused with 15nm -gold (C) 0hr, (D) 4hr post ICV infusion. Subarachnoid space (SAS) of the brain stem following 1.9nm-gold infusion (E) 0hr, (F) 4hr post infusion. The SAS of brainstems following 15nm-gold ICV infusion (G) 0hr or (H) 4hr post. Spinal cords with protruding nerve roots from animals ICV infused with 1.9nm-gold after (I) 0hr or (J) 4hr. Spinal cords with protruding nerve roots from animals ICV infused with 15nm-gold after (K) 0hr or (L) 4hr. Dotted lines mark nerve roots. Arrowheads mark ink cuffs. Trigeminals (M) 0hr or (N) 4hr post infusion of 1.9nm-gold tracer. Trigeminals (O) 0hr or (P) 4hr post 15nm-gold ICV infusion. Dotted line mark ink cuffs. Panels have a minimum N = 3.

We sectioned and performed enhanced gold staining on the 1.9nm and 15nm nanogold infused tissues for more sensitive localization of the infused nanogold (Fig. 3). CNS tissue sections from 1.9nm and 15nm ICV animals with enhanced gold staining showed very similar nanogold distribution at 0hr, as was seen in whole tissues (Fig.3A, C, E, G). At 0hr post infusion both sizes of nanogold were primarily located in the meningeal layers with minimal staining of the parenchyma. By 4hr post infusion 1.9nm nanogold was well distributed throughout the parenchyma of the brain and spinal cord with staining maintained in the meninges (Fig.3 B, F).

**Figure 3.**
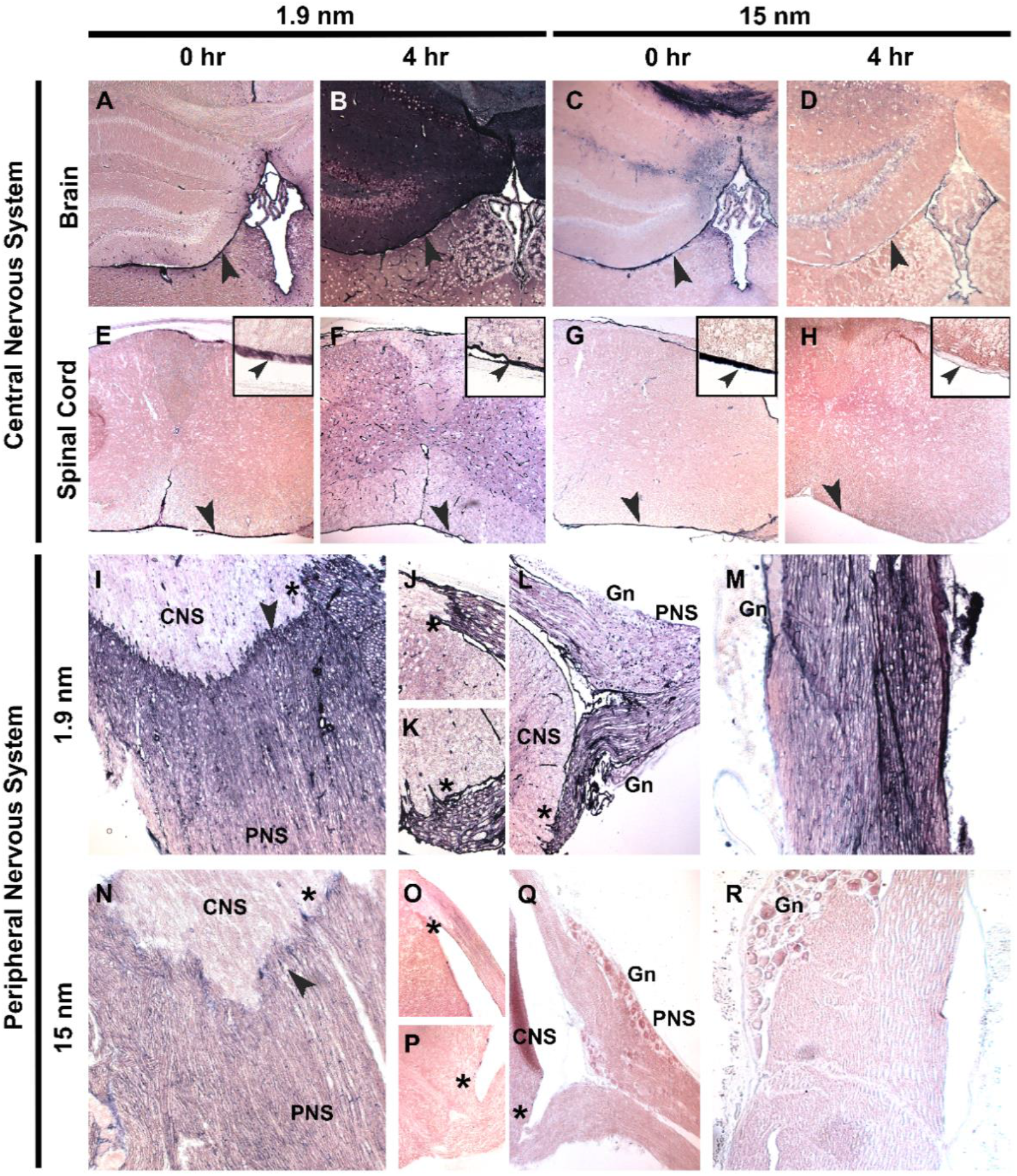
CSF flow dependent on probe size. 1.9 or 15nm nanogold tracers were infused into the lateral cerebral ventricle of fully anesthetized mice. CNS and PNS tissues were harvested 0hr or 4hr after nanogold infusion processed with GoldEnhance. Coronal sections of brain from animals injected with 1.9nm-gold (A) 0hr or (B) 4hr post-infusion. Coronal sections of brain from ICV infused 15nm nanoprobe (C) 0hr or (D) 4hr post-infusion. Arrowheads mark positively staining brain tissue. Spinal cords ICV infusion 1.9nm-gold after (E) 0hr or (F) 4hr spinal cords with nerve roots from animals ICV nfinfusion 15nm-gold(G) 0hr, (H) 4hr post-infusion. Arrowheads mark meninges. “*” marks CNS-to-PNS transition all images. (I) Trigeminal nerve 1.9nm-infused animals. 20X (J) Dorsal nerve root, (K) ventral nerve root from (L). (L) Spinal nerve roots, (M) Sciatic nerve 1.9nm-gold infused animal. Gn = ganglion. (N-R) Peripheral nerves from 4hr 15nm-gold infusion. (N) Trigeminal nerve from 15nm infused animals. High magnification image of (O) dorsal nerve root and (P) ventral nerve root from (Q). (Q) Spinal nerve roots from 15nm-gold infused animals. (R) Sciatic nerve from 15nm-gold ICV infused animals. Panels have a minimum N=4.

Larger 15nm nanogold showed minimal distribution within the parenchyma of the brain and spinal cord with clearing of the meningeal layers for both tissues by 4hr (Fig.3D, H). We next examined peripheral nerves isolated 4hr post infusion with 1.9nm and 15nm nanogold. 1.9nm infused trigeminal, dorsal/ventral nerve roots continuing into the brachial plexus and the sciatic nerve all showed areas of saturated nanogold staining as seen previously (Fig. 3 I-M). Only the trigeminal nerves of 15nm infused animals showed trace amounts of gold staining near the CNS to PNS transition (Fig.3N). Neither the spinal roots nor sciatic nerves of 15nm infused animals showed staining above PBS infused control background at 4hr (Fig. 3O-R, S5). These results indicate size restriction of CSF nanogold distribution in peripheral nerves.

### Nanogold infused into the CSF transits the length of peripheral nerves

Our initial studies demonstrated extensive distribution of 1.9 nm nanogold in the proximal regions of the endoneurium but did not show nanogold distribution throughout an entire peripheral nerve. To increase the sensitivity of detection, we performed injections with 1X and 2X concentrations of nanoprobe in the same 10 μL infusion with PBS infused tissues as controls. In previous gold enhanced staining we stained for one quarter of the manufacturer’s suggested staining time to avoid excessive saturation, allowing for improved morphological assessment. To test for complete distribution of nanogold throughout the trigeminal nerve we enhanced staining for the recommended 20 minutes. Whole trigeminal nerves (from CNS junction at the brain following the maxillary nerve branch to where it innervates the cheek) were assessed from 1X and 2X concentration nanogold infused versus PBS infused control animals (Fig. 4 and Fig. S6 for quantification). Probe deposition within the perineurium traversed the entire length of the trigeminal nerve in both the 1X and 2X. Doubling the concentration of nanoprobe infused into animals doubles the distance of probe deposition within the endoneurium (identified by labeling for the nodes of Ranvier) extending throughout the entire dissected trigeminal nerve (Fig. 4, S6B). In addition, 2X nanogold infused animals had significantly larger “saturated” staining zones defined by maximum number of strong positive pixel values measured by the Aperio ImageScope software. Following the saturated staining was a linear decrease in staining intensity extending to the end of the dissected trigeminal nerves, whereas trigeminals from 1X nanogold infused animals reached background staining levels well before the end of the nerves (Fig. S4A). Collectively, these results indicate visualization of CSF flow is concentration dependent, allowing for identification of nodal labeling to the distal ends of the trigeminal nerve.

**Figure 4.**
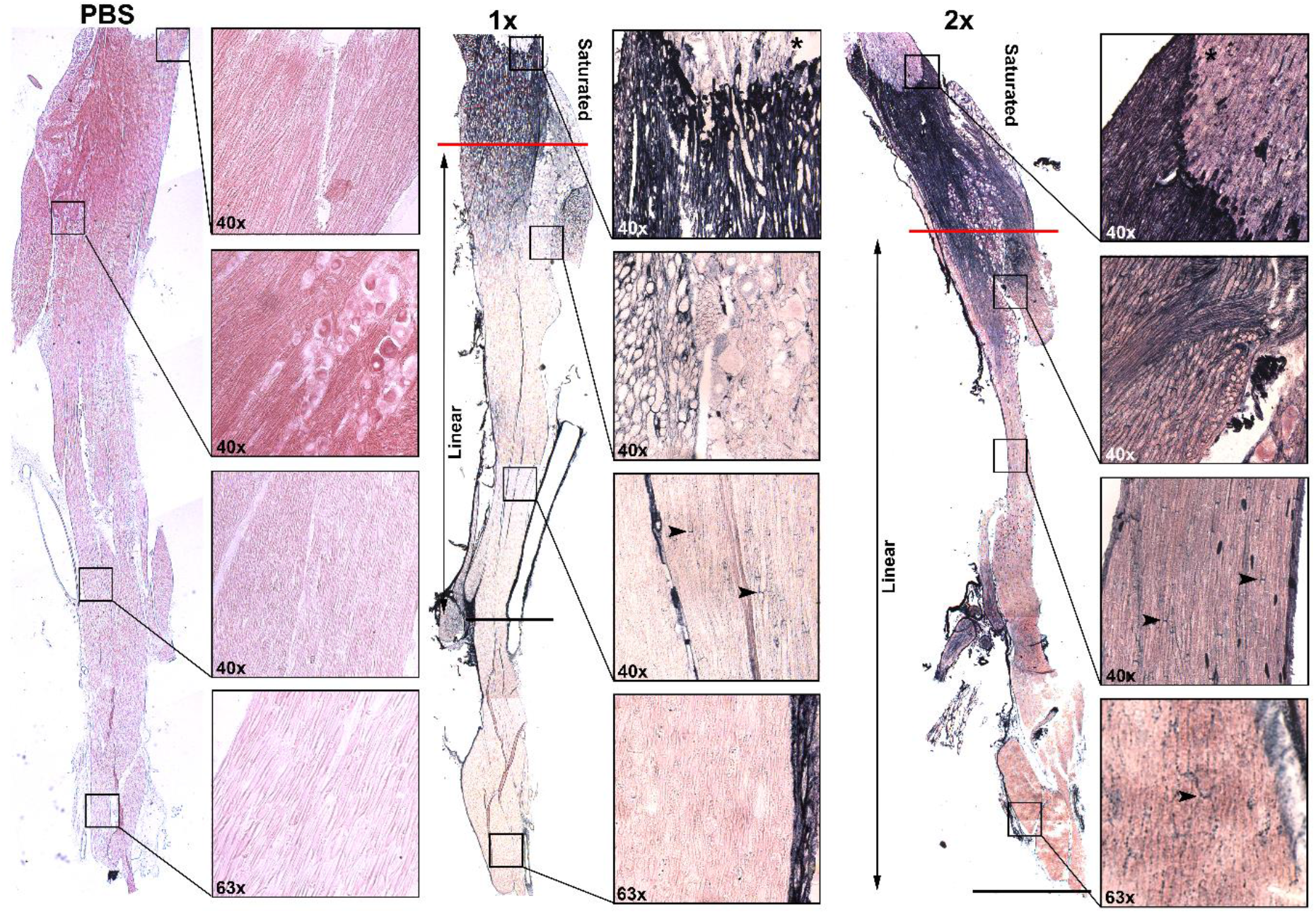
CSF flow of nanogold is concentration dependent. 1x and 2x concentration of nanogold was infused into the lateral cerebral ventricle. Trigeminal nerves were harvested 4hr after infusion and stained with gold enhancement for 20 minutes. 5x images were tiled to show the distance of saturated staining (red line). The distance of CSF infused nanogold flow was determined by measuring the distance for the last nodal labeling seen from the transition marked by a black line. The linear zone marks the decrease in concentration of nanoprobe from the end of the saturated zone to the last nodal labeling seen. For each concentration, 40x and 63x inlays show representative staining at 4 distances. PBS infused nerves only exhibit background staining through the length of the nerve. The saturated zone on 1x concentration nerves reaches approximately 750 μm from the transition, and the last nodal staining was seen on average halfway along the nerve. The saturated zone on 2x concentration nerves was on average 1000 μm from the transition. Nodal staining was seen for the length of the nerve. Panels have a minimum N=4.

### CSF flow from CNS to peripheral nerves is contiguous and continuous

CSF flow within the CNS is known to be a slow, low-pressure system when compared to the vascular system(8, 9, 23). which would suggest CSF driven distribution of nanogold may require significantly longer times than intravenous distribution. Thus, we next sought to determine the effect of time on CSF nanogold flow within peripheral nerves. We performed a time course analysis to determine peak staining intensity for peripheral nerves following a bolus of 1.9nm nanogold at 1X concentration infused into the lateral ventricle with PNS and associated CNS tissues harvested at 0hr, 2, 4, or 6hr post infusion (Fig. 5). Fig. S7 shows pixel intensity quantification of nanogold distribution confirming that increased distance from the infusion site results in decreased maximal staining, requiring increased time for peak staining. The highest concentrations of nanogold were seen within the CNS structures. The brain and spinal meninges label immediately following infusion (Fig. 3 A, E, 5F, S2). However, staining within peripheral nerves varied based on distance from the infusion site (Fig. 5, S7A). The highest staining within peripheral nerves is seen in within the skull, proximal to the injection site, diluting as it flowed distally passed the spinal nerve roots, with the lowest intensity of the nerves assessed seen in the sciatic nerves (Fig.5, S7, S8). Trigeminal nerve staining initiated by 0hr post infusion and peaked at 2-4hr post (Fig. 5B-E, Fig. S7B). The transition from CNS-PNS (measured by 0-200μm from the start of the transition) reached its maximum by 2 hours, while the endoneurial staining (measured by 200-400μm from the start of the transition) did not equilibrate with the transitional labeling until 6hr post infusion (Fig. 5B-E, S7B). Spinal nerve roots began labeling 0hr post infusion albeit at a lower level then the trigeminal (Fig. 5F, Fig. S7A). The most significant change in nerve root staining was seen between 2 and 4hr post infusion (Fig. S7B), exhibiting peak nanogold staining at 4hr post infusion (Fig. 5F-I, Fig. S7). The cervical nerve root and each consecutive nerve root gets a lesser initial concentration leading to a lower overall staining intensity, like that seen in the sciatic nerve (Fig. 5, S7, S8). Fig. S8A-E shows an additional series of cervical, thoracic, and lumbar nerve roots progressing down the spinal cord with additional examples of peak staining times increasing with distance from the infusion site.

**Figure 5.**
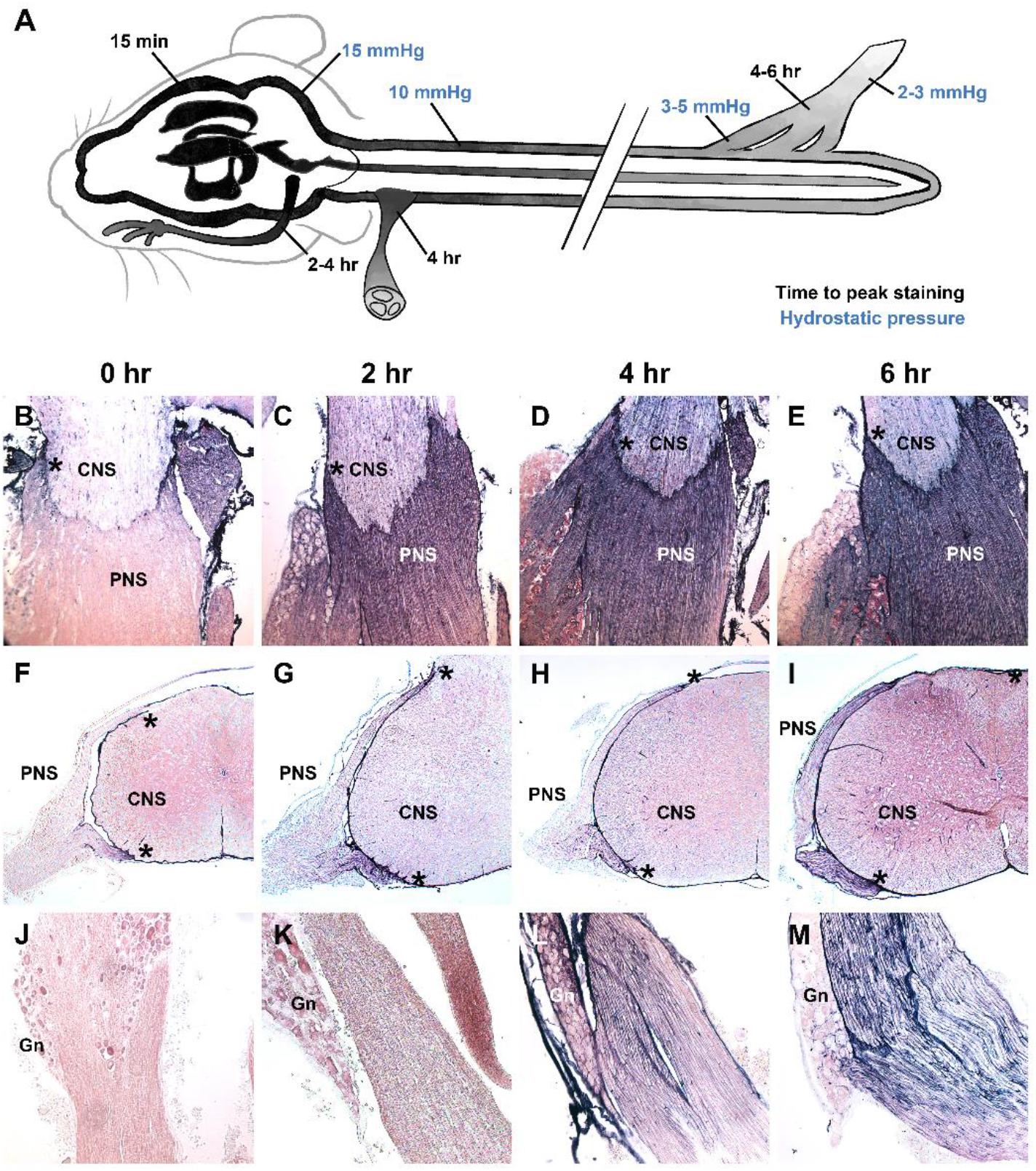
CSF flows through peripheral nerves in a time and distance dependent manner. 1.9nm-gold tracer was infused into cerebral lateral ventricle. CNS and PNS tissues were harvested 0, 2, 4, or 6 hours (hr) post infusion, and then cryosections were stained via GoldEnhanced. (A) Schematic of gold diffusion through central and peripheral nervous system tissues; intensity of black is representative of relative amounts of nanogold and depicts progressive dilution of probe. (B – E) Trigeminal nerves from animals (B) 0hr, (C) 2hr, (D) 4hr, or (E) 6hr post-nanogold ICV infusion. (F – I) cervical spinal nerve roots of 1.9nm-gold ICV infused animals (F) 0hr, (G) 2hr, (H) 4hr, or (I) 6hr post-injection; an “*” marks the transition from CNS to PNS in all panels. (J – M) sciatic nerves of 1.9nm-gold ICV infused animals after (J) 0hr, (K) 2hr, (L) 4hr, or (M) 6hr. Gn = ganglion. Panels have a minimum N=4.

Multiple theories have been advanced for where the CSF would transit into peripheral nerves – the most prominent suggestions being the junction of the subarachnoid angle (SA in Fig. S9A), or the root attachment zone (RAZ in Fig. S9A)(4, 11, 12, 18, 24). Nanogold distribution patterns at the CNS to PNS junction reflect the route of CSF solute transport. Nanogold staining always initiated and extended from the RAZ even though the meninges of the spinal cord were saturated with nanogold (Fig. S9B, C). Staining of distal sciatic nerve also peaked at 4-6hr, similar to the lumbar roots that combine to form the sciatic nerve (Fig. 5J-M, S7C).1X nanogold infusion peaks between 4-6 hours but doubling the concentration (2X) of nanogold infusion sees peak sciatic staining between 2-4hrs (Fig. S8F, G). Peak staining intensity of spinal nerve roots decreased with distance from the infusion site, suggesting continual dilution and efflux of nanogold over time and distance traveled from the infusion site (Fig. 5A).

The junction of CNS to PNS is apparent where the trigeminal nerve (PNS) emerges from the brainstem via the pons (CNS) and where the spinal nerve roots (PNS) branch from the spinal cord (CNS). At CNS/PNS transition, nanogold accumulated at both cranial peripheral nerves and spinal nerves (Fig. S10). At the cranial CNS/PNS transition, increasing gold accumulation over time was indicated by black gold deposits at cells jutting into the CNS, where the demarcation (red line) between CNS and PNS is most clear at 2-6hr post-infusion (Fig. S10a-h). At the junction between peripheral nerve and the brain, nanogold distribution into the peripheral nervous system appears to start at the outside edges of the nerve and fill in more towards the center of the nerve with increasing time; this is especially apparent in comparing trigeminal junctions between 0hr and 2hr tissues (Fig S10A and B, E and F). Endoneurial labeling is seen under the staining at the transition at the 0hr timepoint (Fig. 5B, S10A, E). More distal portions of the peripheral nerve body beyond the initial accumulation point continue to fill over time as the nerve body becomes saturated with nanoprobe (Fig S7B, S10E-H). Probe distribution at the spinal nerve roots reveals the route of probe entry into spinal nerves. Similar patterns of staining to the trigeminal were seen at the spinal CNS/PNS transition (Figure S10I-P). Here, individual axons emerging from the spinal cord parenchyma to the rootlet are darkly stained compared to the adjacent CNS parenchyma in both dorsal and ventral nerve roots. We observe heightened nanogold distribution along cells of the PNS compared to the CNS parenchyma. Quantification of nanogold staining showed a trend for ventral roots to stain more than dorsal roots that was consistent but not significant across all times and samples (Fig. S10 Q).

### Nanogold efflux from the CNS post ICV infusion follows established patterns for CSF outflow

To further validate our CSF probe flow patterns, we evaluated probe deposition in established CSF outflow spaces as recently described by Ma et al(7). Tracers are generally cleared from the CNS through the lymphatic system, which feeds into the cardiovascular system, with ultimate clearance occurring via kidney filtration and concentration in the bladder for excretion(2, 7). We observed ICV infused nanogold distribution followed expected distribution patterns within the CNS (Fig. S2). Nanogold clearance from nervous tissues also followed the expected path in cervical lymph nodes of animals shortly after ICV infusion (Fig. 6A-D), but with significantly delayed kinetics versus IV infused nanogold (Fig. 6E-H). IV infused animals had clear bladders and kidneys within an hour (data not shown), with minimal residual nanogold seen in cervical lymph nodes and spleen by 6hr (Fig. 6E-H). In contrast, animals receiving ICV infusion of nanogold had significant quantities of nanogold in cervical lymph nodes, spleen and kidney, and had black bladders from the concentrated gold after 6hr (Fig. 6A-D). This data shows nanogold is cleared from the nervous system in the same manner as previous CSF efflux studies(2, 7, 10).

**Figure 6.**
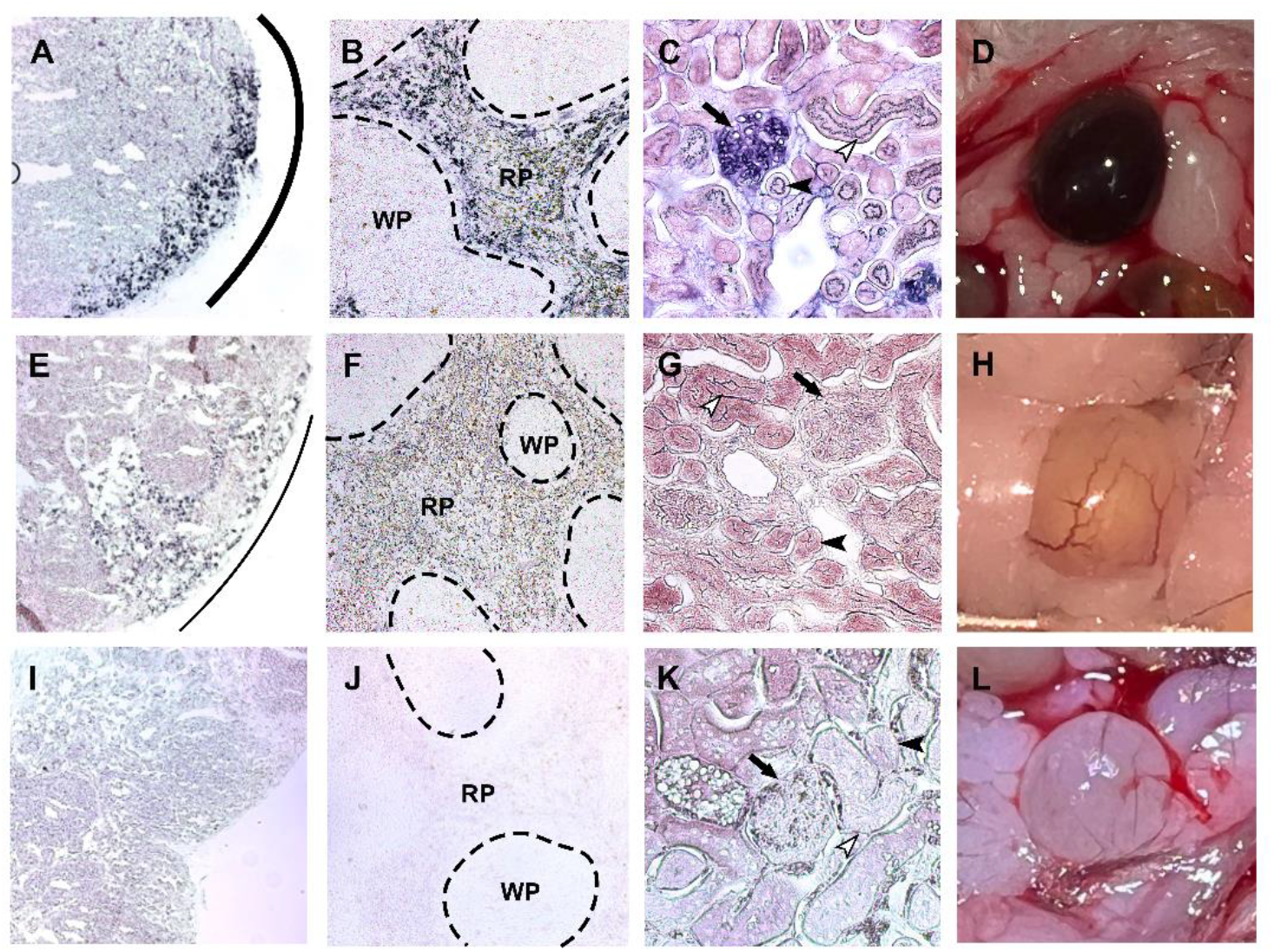
Nanoprobe clearance is delayed in ICV infused nanogold animals. CSF is thought to be effluxed from the central nervous system (CNS) predominately through the lymphatic system with subsequent efflux to the vascular system; final clearance from the body occurs through excretion in the urinary system. To assess clearance of nanogold from the CSF, C57BL/6 mice were ICV infused with 1.9nm-gold tracer (A-D), or intravenously (IV) (E-H) through the tail vein. PBS control ICV infusion and IV injected cohorts (I-L). Tissues were harvested after 6hr, cyrosectioned and GoldEnhanced stained. (A) Lymph nodes, (B) spleen, (C) kidney, and bladder (D) of tissues from ICV-infused animals. (E) Lymph nodes, (F) spleen, (G) kidney, and bladder (H) of tissues from IV-infused animals. (I) Lymph nodes, (J) spleen, (K) kidney, and bladder (L) of tissues from animals infused with PBS into lateral cerebral ventricle.

### Electron microscopy demonstrates CSF infused nanogold delivery to peripheral nerve layers including delivery to axoplasm of peripheral nerves

Gold-enhanced staining for light microscopy generates dark deposits that can obscure histologic detail, making precise identification of labeled cells difficult. Therefore, we used transmission electron microscopy (TEM) for morphological cell identification and ultrastructural localization of nanogold distribution within peripheral nerves (Fig. 7, 8). EM microscopy of peripheral nerve axoplasm showed delivery of nanogold tracer infused into the CSF of brain ventricles, including within axoplasm of distal sciatic nerves (Fig. 8E-J). Nanogold within axoplasm was localized on neurofilaments and within mitochondria of neurons. Perineurial fibroblasts (PF) of the trigeminal nerve contain sufficient nanogold that a minimal enhancement for EM resulted in extremely large deposits along with numerous small to medium deposits within PF (Fig. 7B, C). Lower magnification images show a nearly continual “river” of nanogold intracellularly within the PF, compared to adjacent perineurial cells (Fig. S11A). Perineurial cells did contain nanogold but at low levels compared to fibroblasts (Fig. B7, C and Fig. S11A-C). Analysis of distal lumbar nerve roots (Fig. 7F) and the sciatic nerve (Fig. 7G) showed the same overall staining pattern as trigeminal (Fig. 7E), but with less nanogold found overall. The most obvious difference was the lack of extremely large gold deposits in the PF of the sciatic nerve root, likely due to dilution and efflux of the nanogold bolus as it flows distally (Fig. 5A).

**Figure 7.**
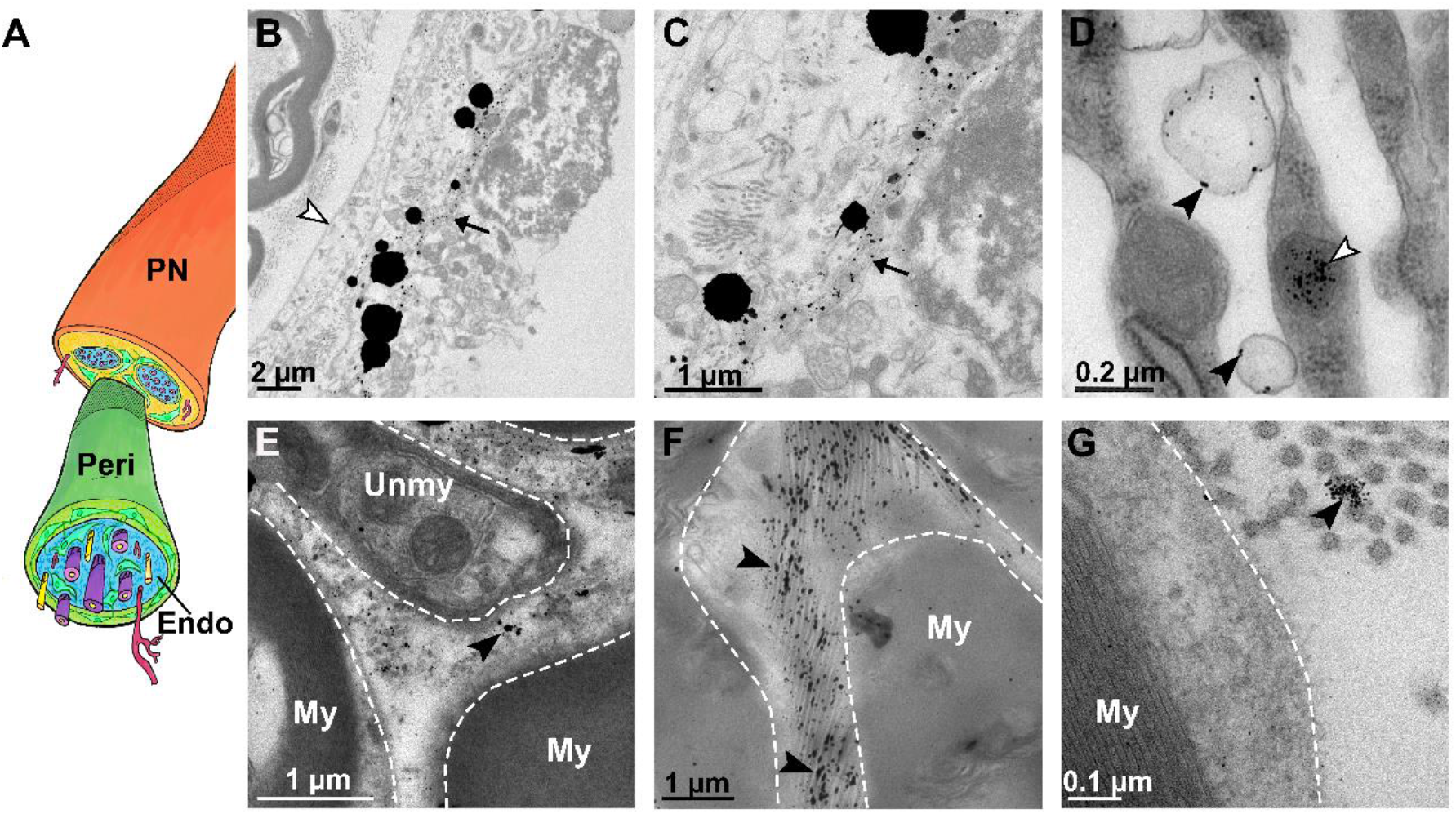
Electron microscopy identifies CSF flows in the connective tissue coverings of peripheral nerves. (A) Schematic of peripheral nerve (PN) that contains multiple fascicles enclosed by the perineurium (Per), which bundles multiple axons (Ax) and their surrounding Schwann cells (Sch); individual axons are located within the endoneurium (En). (B) Cross section of trigeminal nerve 4hr after infusion of 1.9nm-gold tracer into the cerebral lateral ventricle. Nanogold is visible as black particles in fibroblasts (arrow) within the perineurium of the trigeminal after gold enhancement; an adjacent a perineurial cell (open arrowhead) contains little gold. (C) Trigeminal perineurial fibroblast (arrow) with large nanogold deposits. (D) Endoneurial fibroblasts (arrow) in the lumbar nerve root with intracellular nanogold visible following gold enhancement (arrowhead). (E-G) Endoneurial nanogold located in the endoneurial space surrounding myelinated (My) and unmyelinated (Unmy) axons of (E) trigeminal, (F) lumbar nerve root, and (G) sciatic nerves. The endoneurial space is demarcated with dashed lines. Nanogold deposition is seen among collagen fibers in the endoneurium (F, G).

Endoneurium is composed of a loose extracellular matrix (ECM) comprised of collagen fibers bathed in fluid, with a variety of interspersing cells (Fig. 7A). The highest overall levels of nanogold in the endoneurium were in the fluid-filled endoneurial spaces (Fig. 7E-G), with clear deposition in association with the collagen fibrils (Fig. 7F-G and Fig. S13). Tissue and cellular components of endoneurium include small blood vessels, endoneurial fibroblasts (EF), macrophage and mast cells of the immune system, and Schwann cells wrapped around axons. Schwann cells had the most intracellular nanogold (Fig. 8B-D). Myelinating Schwann cells contained more nanogold than did non-myelinating Schwann cells (Fig. S15). As in the perineurium, EF contained abundant nanogold deposits (Fig. 7E-G). Nanogold was also found within endoneurial macrophages (Fig. S16) as well in a variety of intracellular/extracellular vesicles, including extracellular vesicles seen within endoneurial blood vessels (Fig. S17). EM revealed similar nanogold association with ECM structures in the spinal cord as well (Fig. S13). EM of control neural tissues shows no nanogold staining (Fig. S18). Smaller gold deposits were routinely confirmed by electron reflection to differentiate gold particles from normal dark structures (Fig. S11). The EM results agree with the light microscopy staining patterns (Figs. 1-5 and Fig. S19).

**Figure 8.**
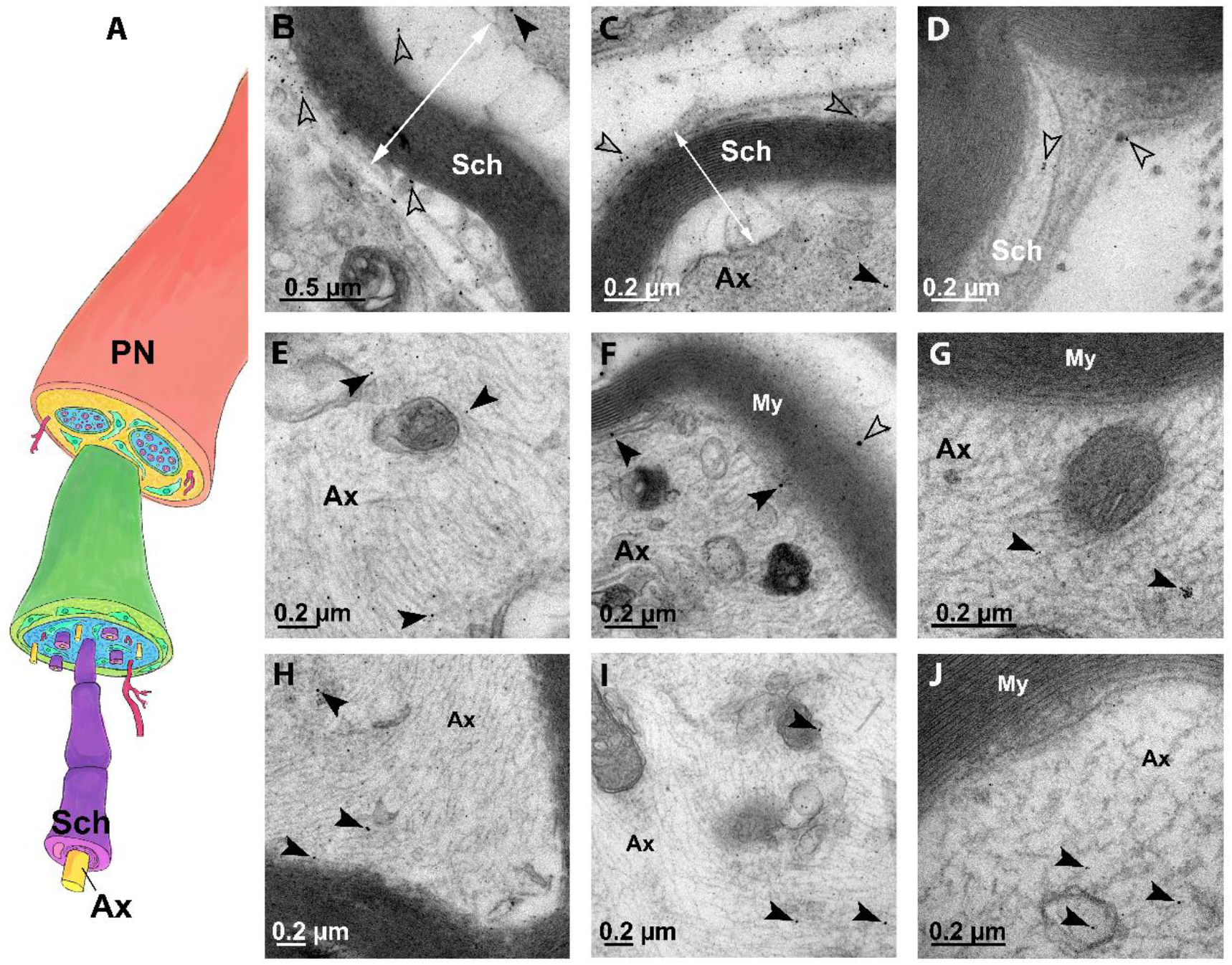
CSF penetrates Schwann cells, down to the axonal level. (A) Schematic of peripheral nerve (PN) that contains multiple fascicles enclosed by the perineurium (Per), which bundles multiple axons (Ax) and their surrounding Schwann cells (Sch). Arrowheads mark examples of 1.9nm-gold tracer in all panels. Examples of nanogold tracer located in the Schwann cell (open arrowheads) and axon (closed arrowheads). (B-D) Nanogold tracer found in the cytoplasm of Schwann cells found in the trigeminal (B, C) and Sciatic nerve (D). (E-J) Nanogold tracer infused into the CSF of mice is found at the Axonal level, located amongst the neurofilament of trigeminal nerves (E, H), lumbar nerve roots (F, I) and sciatic nerves (G, J).

## DISCUSSION

Humans have always been interested in the function of the nervous system. Our minds define our humanity, and our nerves supply the information for our minds. Alterations of the nervous system can affect all systems of the body and all aspects of who we are. Consequently, neuropathies are often reported to contribute to declines in quality of life of patients with numerous diseases. The immune privileged nervous system lacks traditional lymphatics and is protected by the blood brain barrier (BBB), restricting the free movement of molecules(25, 26). In place of blood and lymphatics, cerebrospinal fluid (CSF) functions in the maintenance and health of CNS tissues(16). Bulk CSF flow occurs in the subarachnoid space formed by the CNS meninges (i.e., outer dura mater, middle arachnoid mater, and inner pia mater) supporting its nutrient delivery and waste clearance functions (Fig 1a)(1, 4–6, 12). The CSF flow system contains 150 mL with approximately 500ml of CSF produced per day in humans. CSF is mainly generated by the choroid plexus within the third and fourth ventricles of the brain. CSF then circulates by pulsatile and convective flow to carry nutrients with constant efflux to remove waste and balance production to maintain the low 15mm Hg pressure of the brain (9, 27–30).

Conversely, little is known of the mechanisms maintaining peripheral nerve homeostasis. Nerves of the PNS are surrounded by connective tissue sheathes arranged in an analogous three-layer structure that are continuous with CNS meninges(18, 31). The epineurium is analogous in function to the dura mater(18, 31). The perineurium provides the tight-junction barrier function like the arachnoid mater(32, 33). The endoneurium surrounding axons contains endoneurial fluid (ENF) retained within the “blood-nerve” barrier, a barrier formed by the perineurium and endothelial cells of small penetrating blood vessels of peripheral nerves (17, 34–36). ENF is similar in composition to CSF, and is thought to maintain proper pressure regulation for peripheral nerve health (17). However, the site of production for ENF is not well defined (11–14, 17). The prevailing understanding that CSF flow is restricted to the CNS was established by a variety of vital dye studies using tracers that relied on their inherent properties for detection (not able to be enhanced for more sensitive detection) (1, 4, 5, 22). These studies assessed CSF flow by following tracer distribution patterns. They often described staining “cuffs” forming at the transition from CNS to peripheral nerves with little evidence for tracer flow to peripheral nerves (11–14). Many of these studies have suggested the possibility of CSF flow into peripheral nerves, but the limitations of the vital dyes used, and lack of subcellular resolution did not allow for firm conclusions. These studies provided the basis for the current dogma, inherent in the name cerebrospinal fluid, limiting CSF flow and function within bounds set by the meninges of the brain and spinal cord without significant contribution to PNS function (4, 8, 37).

We recently identified fluid flow channels within nerves of human cadavers which we speculated were possible CSF channels extending from the CNS to peripheral nerves (38–40). In this study we used nanogold particles, infused as a bolus into CSF of the lateral cerebral ventricles, to trace CSF flow patterns in live mice in order to determine if CSF can flow to peripheral nerves under physiological conditions. We selected nanogold to map CSF flow because of several significant advantages as a tracer: 1) It is available in wide range of sizes; 2) nanogold has low osmolality to enhance flow; 3) enhanced gold staining methods allow increased sensitivity of detection by both light and electron microscopy. We selected 1.9nm nanogold, which has an overall particle size of 3.5nm, due to size similarity to proteins typically found in CSF such as albumin (∼3.2-3.8nm) and brain derived neurotrophic factor (BDNF, ∼3.5nm (41–44). Our “large particle” tracer was 15nm nanogold with an overall particle size of 20-30nm which is just above the size of IgG at 15nm, one of the largest proteins found in CSF (45). In contrast, adeno associated virus at 25nm is usually excluded by the BBB/BNB unless leakage is induced, or neurotropic selected serotypes are used (46–48). Recent CSF flow studies using both gold and other nanoparticles have shown restricted flow patterns within the CNS for particles of 20-40nm compared to 3-5-4.5nm, further supporting our size selections (5, 6, 49, 50). Therefore, 1.9nm nanogold would be small enough to potentially transverse the BBB/BNB while 15nm nanogold particles should be mostly excluded. We chose to use the BBB/BNB barrier to guide our selections as it represents one of the tightest barriers in the body resulting in the creation of an immune privileged zone for the nervous system.

The nervous system is traditionally segregated into central and peripheral, leading to research being likewise compartmentalized(8, 51). The current study demonstrates the interconnectedness of the nervous system through the continuity of CSF flow from the CNS to peripheral nerves. This is exemplified by the contiguous distribution of CSF infused nanogold tracer from the lateral ventricle to CNS meninges as well as to PNS perineurium, endoneurium and axoplasm of distal peripheral nerves (diagramed in Fig. 1a). The smaller 1.9nm nanogold traced expected patterns of CSF flow within the central nervous system flowing from the infused ventricles, throughout the brain and spinal cord (Fig. S2) (49). 1.9nm nanogold distribution, however, extended beyond the CNS to the PNS, including distal peripheral nerves (Fig.1,4 and S1). Indeed, by light microscopy, the perineurium exhibits the quickest and highest staining intensity in the peripheral nerve (Fig. 5). We further show ICV infused nanogold flows into the endoneurial spaces of the PNS via light microscopy (Fig. 1, 3-5, S9, S10). High concentrations of nanogold in proximal regions of peripheral nerves obscure morphological details, however, distal segments of nerves with lower nanogold concentrations reveal accumulations of nanogold at the nodes of Ranvier (Fig. 1C, J, K, 4). The nodes of Ranvier are a site for ionic influx required for the generation of action potentials, suggesting the importance of CSF in the propagation of signals along axons along the entire length of peripheral nerves (Fig. 4)(52). Analysis by electron microscopy allowed us to better define the distribution of nanogold within the layers of peripheral nerves. Indeed, within peripheral nerves all cell types examined contained some level of 1.9nm nanogold staining confirmed by the electron reflective properties of gold (Figs. 7, 8, S11, S12). Within the perineurium, fibroblasts contained the highest levels of intracellular nanogold. Enhanced staining of fibroblasts visible as large gold deposits (Fig. S10B, C). The more abundant perineurial cells, however, contain far less intracellular nanogold (Fig. S11) with a concentration pattern similar to CNS tissues (Fig. S14). The axoplasm of peripheral nerves, including axons of the distal sciatic nerve, contained nanogold delivered from the ventricular infusion of CSF within the brain (Figs. 7,8).

Before flowing into the endoneurium, nanogold is seen accumulating at the transition from CNS to PNS as early as the 0hr timepoint in the trigeminal nerve (Fig. 5, S10). The transition of CNS to PNS is demarcated by a clearly visible interface. In both the trigeminal and spinal nerve roots, enhanced gold staining shows less gold staining in CNS tissue with a distinct transition to more highly stained PNS tissue with individual cellular staining clearly visible along the transition (Figs. 1, 3-5, S9, S10). Axons are continuous from the CNS to PNS throughout this area, however a key change happening is in the type of myelinating cell (51). Changes in axon myelination from oligodendrocytes (CNS) to Schwann cells (PNS) occur, even though the axon projection itself can be contiguous across the transition(51). Overall, electron microscopy shows the endoneurium exhibiting the most abundant extracellular nanogold, specifically in the loose collagen fiber matrix where endoneurial fluid is found (Fig. 7E-G). The presence of nanogold in the free-flowing spaces of the endoneurium suggests contiguous CSF fluid flow from the SAS of the CNS to the endoneurium of peripheral nerves. EM also showed the high concentrations of intracellular nanogold within the cytoplasm of endoneurial fibroblasts (Fig. 7D) and Schwann cells (Fig. 8B-D). The high concentration of nanogold found in the fluid space, fibroblasts of the endoneurium and myelinating Schwann cells helps explain the visible enhanced gold staining of the endoneurium with visible nodes of Ranvier seen with light microscopy (Figs. 1, 4, S10). Schwann cells likely aid in nutrient delivery to axons from the endoneurial space gaining access to the axons through the nodes of Ranvier and Schmidt Lanterman clefts, resulting in gold being easily detectable within the axoplasm of peripheral axons (Fig. 8E-J) (53, 54). Nanogold deposition within the axoplasm of peripheral axons suggests that CSF flow in the periphery may be integral for peripheral axon maintenance.

Nanogold delivery to the PNS was time and distance dependent (Fig. 5). Our results demonstrating CSF flow into peripheral nerves using nanogold tracing are diagramed in Fig. 5a. CNS structures label the most intensely the quickest. (Fig. 2, 3, S2). However, the distribution of nanogold to the PNS is a slower process, with staining dependent on distances from infusion site (Fig. 5, S7, S8. S10). Quantification of enhanced gold staining of peripheral nerves showed more proximal to the site of ventricle infusion achieved maximal staining faster with larger zones of saturated staining. The trigeminal nerve was the most proximal peripheral nerve sampled which achieved maximal staining in as little as 2 hours and retained sufficient nanogold to remain at peak saturation through 6 hours post infusion. More distal spinal and sciatic nerves took longer to initiate staining while achieving an overall lower peak staining which peaked around 4hrs and showed evidence of washout by 6hrs post infusion (Fig. S7).

Previous studies identified the accumulation of dyes and tracers at the subarachnoid angle, a point where the arachnoid mater folds back on itself and becoming continuous with the dura creating a “cul-de-sac” where peripheral nerves branch exit from the CNS (4, 11, 12, 14). Our results reveal 1.9nm nanogold infused into CSF entering the PNS through the transition/root attachment zone (RAZ) (Fig 1-5, S9, S10). At this site, nanogold is seen to flow in from the edges of the nerve, suggesting continuity between the transition and the subarachnoid space (Fig. 5, S10). Loading at the transition appears to be required for flow into the endoneurium, as staining in the endoneurium is only seen under a labeled transition Fig. S10A-E). Similar to what was described by Brierley, larger, 15 nm nanogold formed a “cuff” at the transition in both the trigeminal and spinal nerve roots, but was unable to enter the PNS in any significant amount (Fig. 2K,L,O,P, 4)(12). Unlike previous studies however, the small size of the 1.9nm nanogold allows entry into the PNS (Fig 1-8). The restriction of larger particles and flow mechanisms into the peripheral nervous system may be similar to the size restrictions reported for the BBB/BNB and glymphatic systems (5, 55, 56). Further exploration of the size restrictions and mechanisms of CSF flow into peripheral nerves could yield interesting new developments in drug development for nervous system disorders.

The endoneurium is bounded within the perineurium and consists of loose fluid filled connective tissue containing Schwann cells and axons of peripheral nerves. Such a structure would help explain the slow time and size-dependent process of CSF infused nanogold distribution in peripheral nerves (Fig. 2, 3, 5, S10). Although the precise mode of peripheral nanogold transport remains to be determined, it is highly probable that peripheral CSF flow is driven by both convective and pulsatile fluid flow, a model mirrored by what is known of central CSF flow(4, 9, 10, 23). The hydrostatic gradient from CNS to PNS likely drives convective fluid flow from the subarachnoid space of the CNS to the endoneurium along peripheral nerves(17, 57). Thus, the endoneurium may be functionally analogous to the subarachnoid space of the CNS in supporting CSF flow through peripheral nerves. Additionally, the close proximity of peripheral nerves to blood vessels suggests that peripheral CSF flow may be driven by pulsatile fluid flow. Such a model agrees with our knowledge of endoneurial fluid within peripheral nerves, the barrier properties of the perineurium, and the physical continuity of connective tissue coverings between CNS and PNS(18, 31).

The presence of a contiguous CSF flow route from the CNS along peripheral nerves has far-reaching implications for peripheral nerve health and neuropathologies. In the CNS, CSF is known to provide cells with a variety of molecules involved in neural metabolism and maintenance(58). CSF is also known to play a critical role in eliminating waste from neuronal cells(5, 7). We expect that a whole nervous system CSF flow system allows CSF to function with similar roles for peripheral neural tissues, i.e. delivering nutrients and clearing waste and physical cushioning to peripheral nerves. Alterations to normal CSF flow patterns from the CNS to the periphery could result in dysregulated and deficient nutrient delivery and waste clearance functions in peripheral nerves, resulting in aberrant peripheral nerve function. Aberrant CSF flow through the CNS results in accumulation of toxic waste products like neuritic plaques implicated in neuronal diseases(5, 7). In the system we describe here, CSF is continuously produced primarily within the choroid plexus of brain ventricles with subsequent contiguous central to peripheral CSF flow(2, 4). Similar to CNS CSF flow and unlike the high pressure closed circulatory loop of the vascular system, the newly described CSF flow system functions as a low-pressure system with continuous drainage. Under normal conditions, various routes of efflux from the CSF system facilitate drainage and help maintain a low-pressure system. Continuous CSF efflux from peripheral nerves would also enable waste clearance to maintain nerve health. We documented nanogold efflux to the lymphatic and vascular systems with filtration by kidneys and elimination via urine in the bladder (Fig. 6).

Our results suggest that infusion into the CSF compartment may be an efficacious method for direct delivery of drugs and therapeutic agents to peripheral nerves. In the CNS, CSF drainage is facilitated predominately by lymphatic drainage (7). Along cranial peripheral nerves and spinal nerve roots are lymphatic vessels that may facilitate additional CSF drainage from the PNS (2, 4, 27). Following ICV infusion, CSF outflow from peripheral nerves is continuous throughout the 6 hr timepoint, while IV infusions are cleared by 1 hr post infusion (Fig. 6). Further, agents delivered via the cardiovascular system are spread diffusely throughout many tissues and are not delivered to peripheral neural tissues in any appreciable amount (Fig. 1E, I, M) (17, 36). We found that in IV infused animals, nanogold was never detectable beyond the barrier of the perineurium, whereas ICV infusion resulted in direct delivery to the endoneurium of peripheral nerves down to peripheral axons (Fig. 1E, I, M, 10 E-J). Therefore, we expect the development of drugs capable of spanning the central to peripheral nervous system transition through the observed CSF flow system could be a clinically viable route for slow-release drug delivery to peripheral nerves. This direct PNS access could allow for effective doses to be achieved without significant systemic exposure. In addition, gold nanoparticles have shown great potential as therapeutic agents due to their unique physical and chemical properties(59, 60). By showing gold nanoparticles infused into the CSF reaching distal peripheral axons in a nervous system specific manner, we have unlocked a new mechanism for drug delivery targeting central and peripheral nervous system disorders.

Using nanoprobe technology, we provide evidence that CSF transport extends beyond the CNS into the PNS, where it is present within the individual axons of peripheral nerves. Our kinetic results demonstrate a unique continuous flow system of the CNS and PNS that may not only explain some neurological disorders but may also provide an avenue for delivery of therapeutic agents and novel diagnostic imaging for peripheral neural tissues. The identification of central CSF influence on the PNS, and more specifically peripheral axons indicates an interconnectedness of the nervous system that was previously unidentified, a discovery that could revolutionize the way we view and treat nervous disorders.

## MATERIALS AND METHODS

### Animals

C57BL/6 (6 weeks – 5 months) were acquired from Charles River and Jackson Laboratory. CD1^+^ mice (5 weeks – 12 weeks) were obtained from the University of Florida breeding facility and maintained in the AAALAC-inspected UF rodent vivarium. Injections were performed in both inbred (C57BL/6) and outbred (CD1^+^) strains to assess potential variation between strains. Both male and female mice were used to assess variations due to gender; mice were randomly assigned to experimental and control groups. No appreciable differences in nanogold distribution were seen in either strain or sex. All cohorts had a minimum of 4 animals. For nanogold infused animals the success of the infusion was confirmed by histology of the injection site. Animals where the infusion was not within the lateral cerebral ventricle were excluded from test cohorts. All procedures and procedural spaces were approved by the University of Florida’s Institutional Animal Care and Use Committee (IACUC).

### Pre- and post-procedural care

Mice were given pre- and post-procedural analgesia to mitigate pain. Mice were anesthetized, and sufficient depth of anesthetic plane was ensured by loss of foot-pedal reflex for all procedures. Procedural sites were denuded by shaving and disinfected by triple scrubbing with iodine and 70% ethanol. All procedural surfaces were sterilized with an approved sterilant. All surgical instruments were sterilized by autoclave prior to use. Mice were allowed to recover from anesthesia on a heated surface to maintain physiological body temperature. Recent reports have indicated CSF production rates and subsequent flow rates are influenced based on whether animals are anesthetized and differ with different anesthetics(*21, 61*). We observe CNS to PNS flow in both fully anesthetized and awake mice. In our experiments, mice at 0hr time point (harvested directly following ICV injection) exhibit the beginnings of flow spanning CNS-PNS in the trigeminal nerve. Mice at all other time points (2, 4, 6hr) were allowed to fully recover from anesthesia.

### Surgical procedures (injections)

#### Ventricular CSF injections

To assess CSF flow patterns, nanogold (Aurovist^TM^, Nanoprobes Inc., Yaphank, NY) was injected into the lateral cerebral ventricle of C57BL/6 and CD1^+^ mice fully anesthetized with Avertin (250 mg/kg, intraperitoneal (IP) delivery). To access the cerebral lateral ventricle, a mid-sagittal skin incision was made to expose the bregma. A 0.1 mm bur-hole was made into the skull and a 32-gauge canula was then inserted and immobilized in the right ventricle at the following coordinates relative to the bregma: 1 mm lateral, 0 mm antero-posterior, and 2 mm subcranial. 1.9nm Aurovist (0.1 mg/μL), 15nm Aurovist (0.1 mg/μL) or 1x PBS was then continuously infused into the lateral cerebral ventricle using a syringe pump (World Precision Instruments, Sarasota, FL) via a Hamilton syringe (Hamilton Company, Reno, NV) connected to the canula with polyethylene tubing at a rate of 1 µL/min. For standard injections (and unless otherwise specified) injections were carried out over 10 min for delivering a total of 1 mg of Aurovist/nanogold (1/40 the manufacturer’s suggested amount for IV administration). For low volume injections, 2.5 uL of 0.4 mg/μL Aurovist (1.9 nm) was delivered over 2.5 min for delivery of 1 mg nanogold at a flow rate of 1 μL/min.

For negative controls, central and peripheral neural tissues were harvested from either 1xPBS-infused animals or from non-injected animals C57BL/6 or CD1^+^ mice. For PBS-injected control cohort, 1XPBS was infused into the lateral ventricle of anesthetized mice as described above at flow rate of 1 µL/min; mice were euthanized at 15 min, 2hr, 4hr or 6 hr post-injection.

#### Vascular injections

To assess the contribution of the vascular flow to nanoprobe deposition in neural tissues, a 31-gauge needle was used to perform a bolus injection of 1 mg of nanogold in the tail vein of fully anesthetized mice.

Following both cerebral ventricular and vascular injections, animals were euthanized by overdose of Avertin (500 – 800 mg/kg, IP route) followed by intracardiac perfusion with 1X PBS followed by 4% paraformaldehyde (PFA) in 1X PBS.

### Histology and microscopic analysis

#### Tissue harvest and processing

For all animals, brains, cranial nerves, spinal cords (with nerve roots), and peripheral nerves (trigeminal and sciatic nerves) were harvested directly following intracardiac perfusion. Tissues were then drop-fixed overnight at 4°C in either 4% PFA in 1X PBS or in Trump’s fixative for light microscopy or electron microscopy, respectively.

#### Light microscopy

For light microscopy, after 4% PFA fixation, tissues were cryopreserved by sucrose gradient to 30% sucrose (in 1X PBS) and then embedded and frozen in Tissue-Tek O.C.T. Compound (Sakura, Torrance, CA). All tissues were obtained at 5 um thickness using a Leica CM 1859 cryostat (Leica, Germany) and stored at -20°C until use. To assess nanogold deposition patterns, neural tissue sections were “enhanced” with GoldEnhance LM (Nanoprobe, Yaphank, NY) following manufacturer’s instruction. Tissues from both 1.9 nm and 15 nm nanoprobe experiments were developed with GoldEnhance LM following manufacturer’s recommendations. Unless otherwise indicated, tissues were developed with the enhancing solution for 5 min. Tissue sections were coverslipped with Vectashield Antifade Mounting Media (Vector Laboratories, Newark, CA). Sections were analyzed via light microscopy (Leica DM5500 B, Leica Microsystems, Germany). Images were captured with Hamamatsu C7780 camera (Hamamatsa, Japan). Enhanced tissue sections were analyzed by a neuropathologist for blinded review.

#### Electron microscopy

For electron microscopy analysis, harvested tissues were post-fixed by immersion in Trump’s Fixative (containing 4% paraformaldehyde, 2.5% glutaraldehyde in 0.1M sodium cacodylate containing 2mM MgCl2, 1mM CaCl2 0.25% NaCl, pH 7.24) (Electron Microscopy Sciences, Hatfield, PA) overnight at 4°C. Fixed tissue was processed with the aid of a Pelco BioWave Pro laboratory microwave (Ted Pella, Redding, CA, USA). Samples were washed in 0.1 M sodium cacodylate (pH 7.24), post-fixed with buffered 2% OsO4 over night at 4°C, water-washed and then dehydrated in a graded ethanol series (25% through 100% with 5-10% increments) followed by 100% anhydrous acetone. Dehydrated samples were infiltrated with ARALDITE/Embed (EMS, Hatfield, PA) and Z6040 embedding primer (EMS, Hatfield, PA) in increments of 3:1, 1:1, 1:3 anhydrous acetone:ARALDITE/Embed followed by 100% ARALDITE/Embed. Semi-thick sections (500 nm) were stained with toluidine blue. Ultra-thin sections were collected on carbon coated Formvar 100 mesh grid (EMS, Hatfield, PA) and post-stained with 2% aqueous uranyl acetate and lead citrate (EMS, Hatfield, PA). Sections were processed with GoldEnhance EM Plus^TM^ (Nanoprobe, Yaphank, NY) following manufacturer’s instructions to enhance nanogold. Sections were examined with a FEI Tecnai G2 Spirit Twin TEM (FEI Corp., Hillsboro, OR), and digital images were acquired with a Gatan UltraScan 2k x 2k camera and Digital Micrograph software (Gatan Inc., Pleasanton, CA). Electron reflection was used to confirm the presence of gold versus non-reflective native structures. Developed tissue sections were analyzed by a neuropathologist for blinded review.

### Statistical Analysis and Data Preparation

A minimum of 3 biological and 3 technical replicates were used for trigeminal nerves, cervical spinal nerve roots, and sciatic nerves at each timepoint of 0hr, 2hr, 4hr, and 6hr. Cryosections were imaged for quantification following gold enhanced staining with a 5-minute enhancement time, as previously described. Distances of 0-200 μm and 200-400 μm from the transition were outlined using the Aperio ImageScope software (Leica Biosystems). 0-200 μm was used to denote staining at the transition, whereas the distance of 200-400 μm was used to assess endoneurial staining proximal to the transition. Then, the percent positive staining for each region was computed using the Aperio Positive Pixel Count program (Aperio, Vista, CA, USA). The Positive Pixel count program was configured to detect the purple/black color from the enhancement stain (ImageScope, Aperio Technologies, Vista, CA, USA). The total number of purple/black pixels (positive) divided by the total number of all pixels (positive + negative) was represented as a % positive staining for all tissues, the % Positive staining values of all replicates (biological and technical) at each timepoint were averaged by distance or by the distances combined. For the analysis of saturated staining, the total number of strong positive pixels was divided by the total number of pixels (positive + negative). Unpaired t-test was performed to determine statistical significance (* P < 0.05), ** P < 0.01, *** P < 0.001, **** P < 0.0001).

All statistical and graphical representations were created using Graph Pad Prism version 9. Final images were assembled using Adobe Photoshop.

### List of Supplementary Materials

Fig. S1 S19

S1 Low volume, equal concentration nanoprobe injections yield same nanoprobe deposition patterns as standard injection

S2 Nanoprobe distribution recapitulates known CSF flow routes in central nervous system (CNS).

S3 Average nanogold staining of peripheral nervous system compared to the central nervous system.

S4 1x PBS control of whole, unenhanced tissues.

S5 Control 1x PBS histological sections following gold enhance staining.

S6 Gold enhanced stain intensity and distance quantification of 1x and 2x concentration.

S7 Average nanogold staining of peripheral nervous tissue after 0, 2, 4 and 6 hours.

S8 CSF flows through spinal nerves in a time and distance dependent manner.

S9 CSF solute enters PNS at nerve root attachment zone.

S10 CSF solute accumulates in PNS at CNS/PNS junction.

S11 Perineurial cells contain lower levels of nanogold equivalent to CNS levels of staining.

S12 Electron reflection verifies nanogold by electron microscopy.

S13 Nanogold accumulates on collagen fibers in the peripheral nervous system (PNS).

S14 Electron microscopy of central nervous system tissue.

S15 Unmyelinated Schwann cells exhibit lower levels of intracellular nanogold.

S16 Nanogold is phagocytosed in peripheral nerves.

S17 Nanogold accumulates in extracellular and intracellular vesicles.

S18 Electron microscopy of trigeminal nerves from control animals.

S19 Light Microscopy of the nerves used for electron microscopy.

## Acknowledgments

The authors thank Rudy Alvarado and Nicole Machi of the University of Florida, Interdisciplinary Center for Biotechnology Research, Electron Microscopy Core. This work was supported by RO1 DK 121117 to EWS and T32 DK074367 to EWS, APL and AVV.

## Author contributions

APL and AVV are co-first authors and contributed equally. All authors made critical comments related to the intellectual content of the manuscript.

Experimental Design: APL, AVV, JEP, and EWS

Methodology: APL, AVV, JEP, JAC, DEB, and OGF

Blinded reviewers: ATY and WAD

Data analysis: APL, AVV, JEP, ATY, WAD, DES, and EWS

Diagram design: EAS

Supervision: EWS

Funding acquisition: EWS

Writing – original draft: APL, AVV, and EWS

Writing – review & editing: APL, AVV, ATY, WAD, GAJB, EWS

## Data and materials availability

All data needed to evaluate the conclusions in the paper are present in the paper and/or the supplementary material.

## Supplemental Figures for “Cerebrospinal Fluid Flow Extends to Peripheral Nerves”

**Fig. S1.**
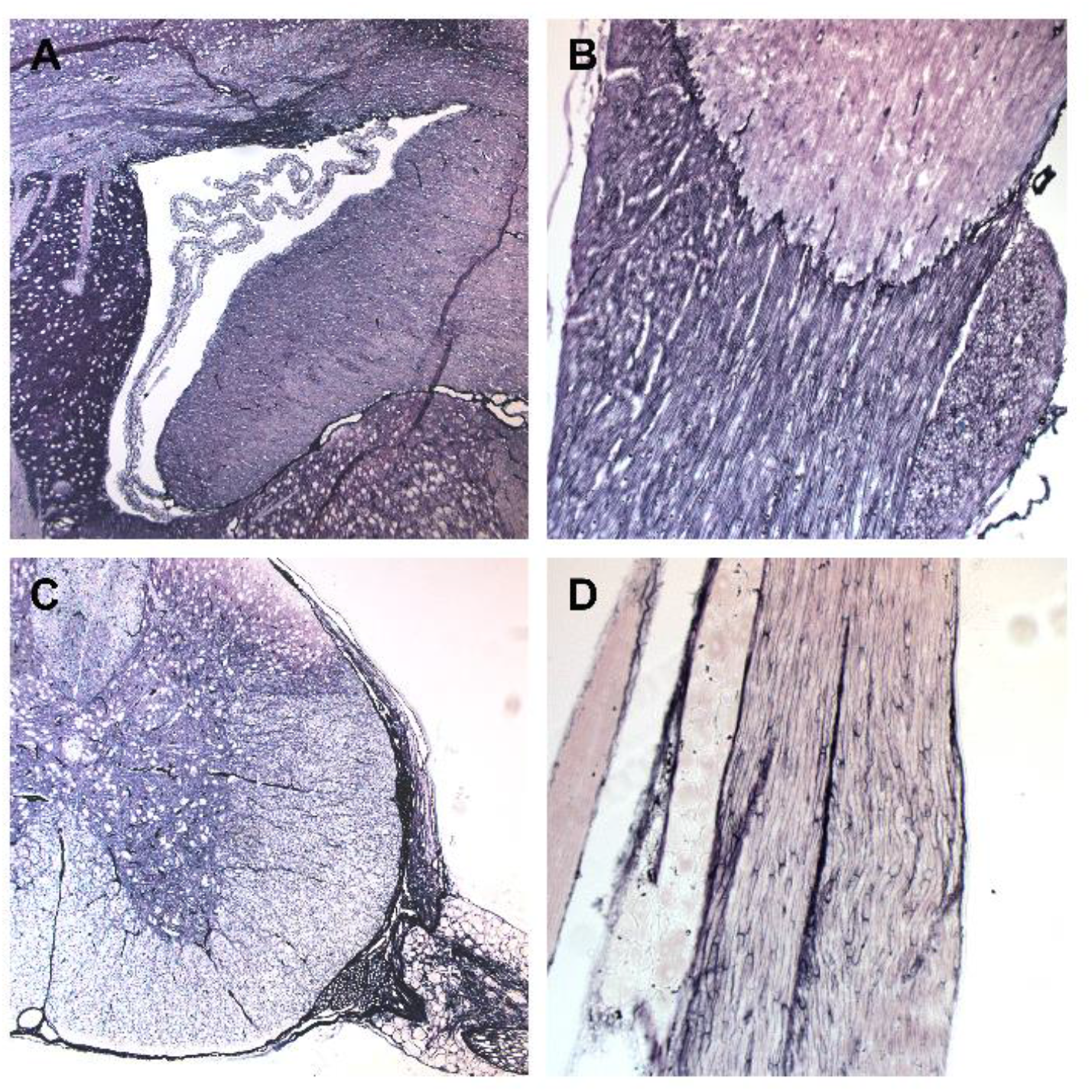
Low volume, equal concentration nanoprobe injections yield same nanoprobe deposition patterns as standard injection. To control for the injection volume used, fully anesthetized C57BL/6 mice were injected with 0.4 mg/μl 1.9 nm nanogold tracer in a 2.5 μl aliquot over 2.5 min (1 μl/min) into the lateral cerebral ventricle. Central and peripheral nervous system tissues were harvested at 4 hours post injection. Cryosections were developed with GoldEnhance using standard protocol. (A) Lateral ventricular (LV) injetion site. (B) Coronal section of spinal cord. (C) Trigeminal nerve. (D) Sciatic nerve. Comparable staining of central and peripheral nervous system tissues were seen, however, the small volumes of more concentrated and more viscous nanoprobe led to concerns of a varied amount of probe delivery. N=4 for all panels.

**Fig. S2.**
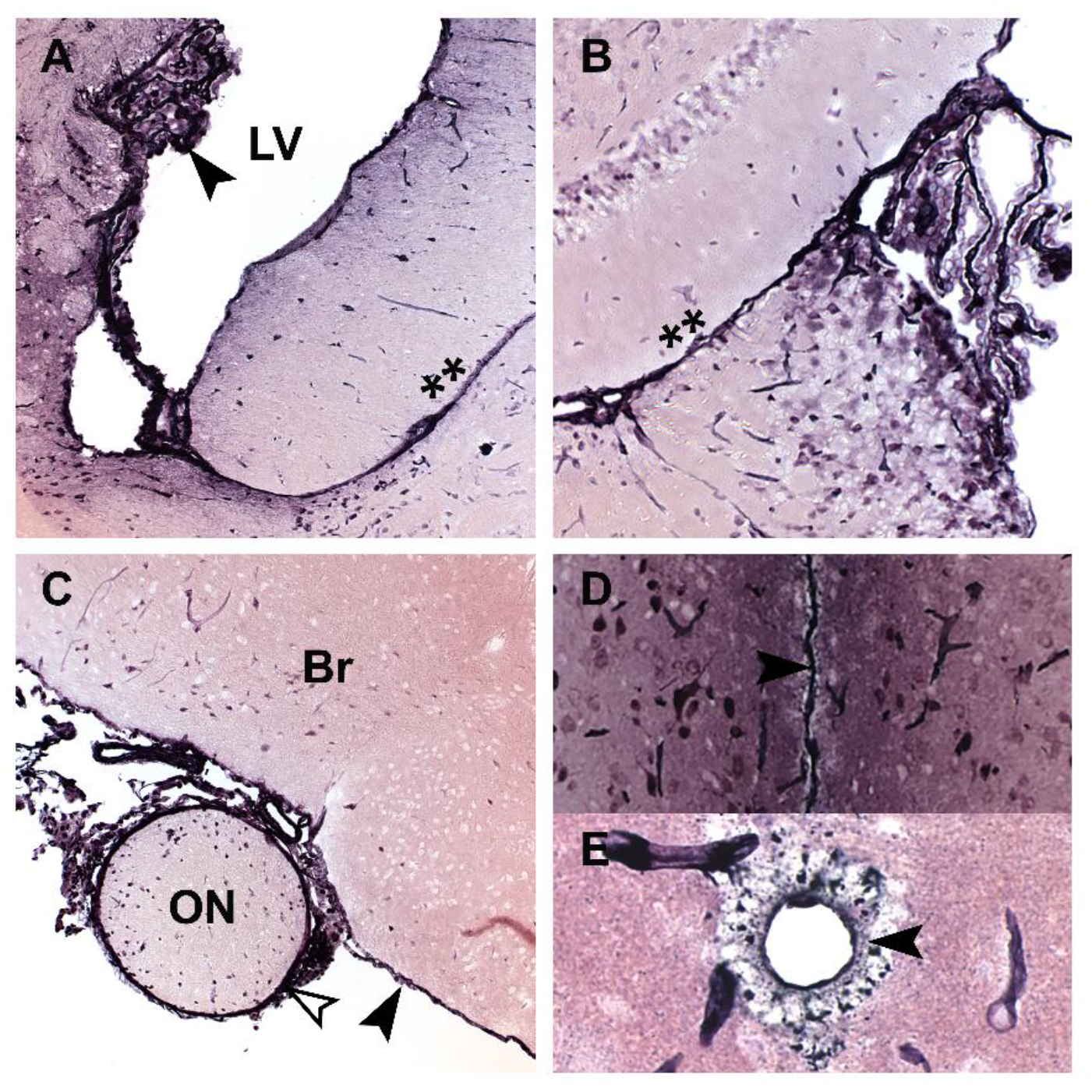
Nanoprobe distribution recapitulates known CSF flow routes in central nervous system (CNS). To validate our CSF tracing system, we sought to assess nanoprobe distribution patterns in the central nervous system (CNS) directly after intracranial injection of 1.9 nm nanogold tracer into fully anesthetized mice. Tissues corresponding to established CSF flow sites were harvested from animals 15 min after nanoprobe infusion. Cryosections were obtained and enhanced with GoldEnhance LM for 5 min. Coronal brain sections depicting lateral ventricular (LV) infusion site (A), choroid plexus (B), meninges of brain (Br) and optic nerve (ON) (C), cerebral aqueduct (D) and central canal (E). These results reveal our particle tracing system faithfully recapitulates CSF flow in the CNS and helps to validate our CSF tracing system for the peripheral nervous system.

**Fig. S3.**
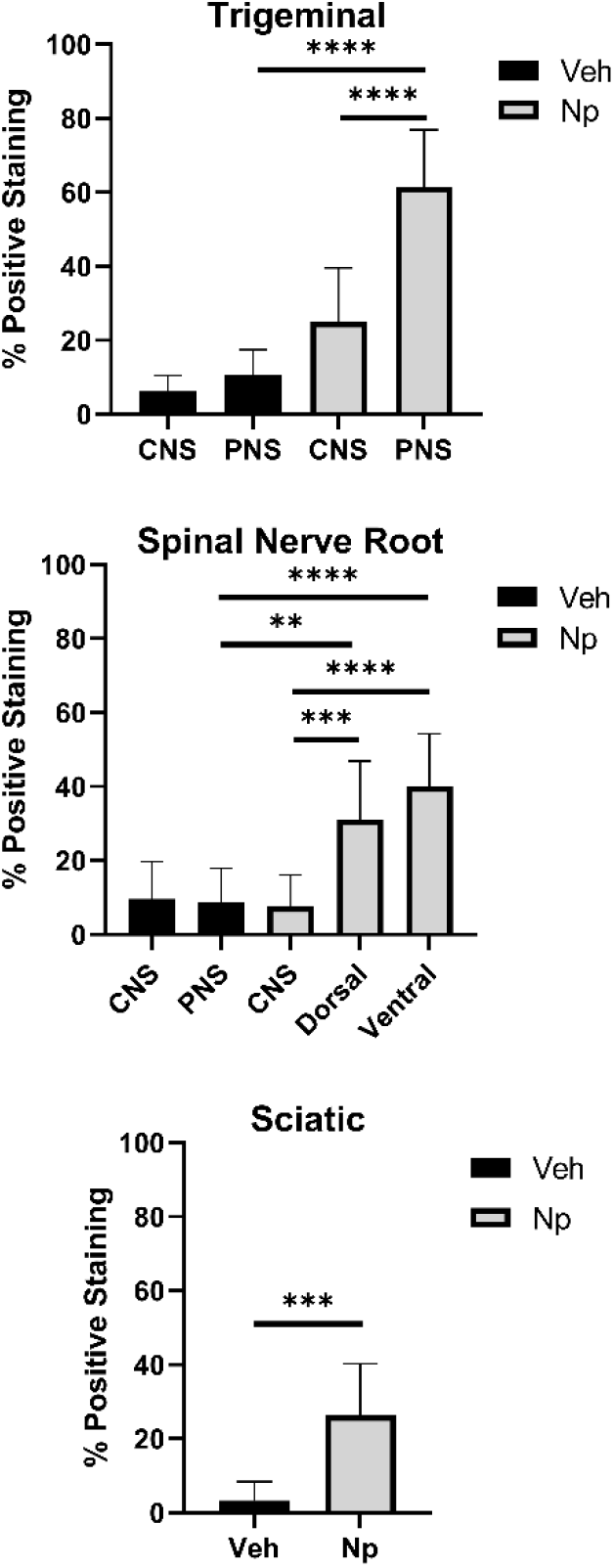
Average nanogold staining of peripheral nervous system compared to the central nervous system. To elucidate probe deposition patterns in 1.9 nm nanogold tracer ICV-infused mice, we quantified positive pixel density. C57BL/6 mice were injected with 1.9 nm Aurovist into the lateral cerebral ventricle, and trigeminal, cervical spinal cord, and sciatic nerve tissues were collected 6 hr after probe infusion. Tissue samples were cryosectioned and developed with GoldEnhance for 5 min. To determine probe abundance compared to PBS-infused animals, positive pixel density was measured in the CNS and PNS of trigeminal nerves, spinal nerve roots, and sciatic nerves 6 hrs after 1.9 nm nanogold (Np) or 1x PBS infusion (Veh). Positive staining was compared between CNS and PNS in Veh and NP infused mice for the trigeminal and spinal nerve root. Additionally, dorsal and ventral spinal nerve roots were compared. Sciatic nerves were compared between Veh and Np. PNS staining was statistically significant compared to both the CNS of 1.9 nm nanoprobe infused and CNS/PNS of 1x PBS infused tissues. Unpaired t-test was performed to determine statistical significance (** P < 0.01, *** P < 0.001, **** P < 0.0001).

**Fig. S4.**
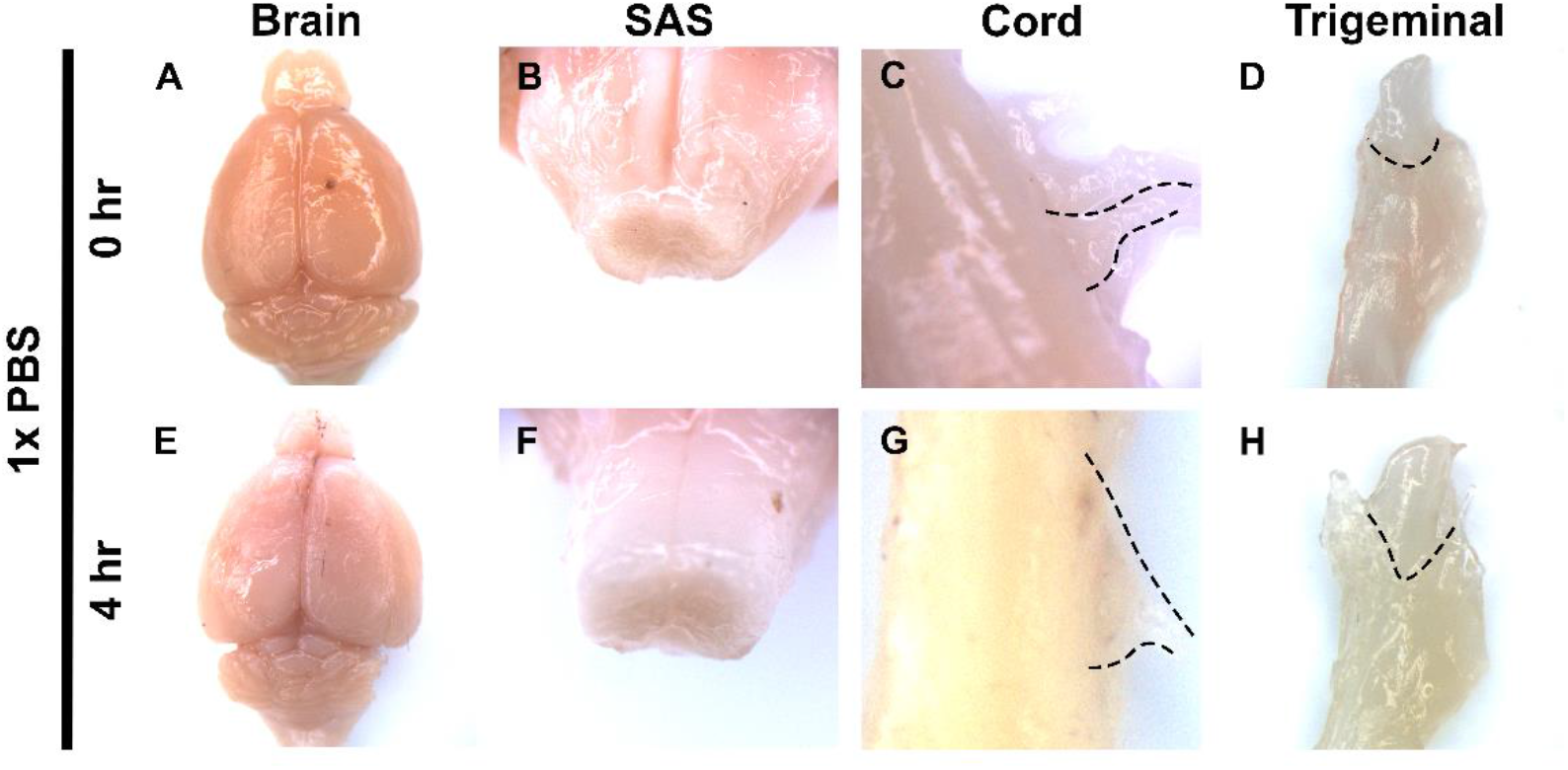
1x PBS control of whole, unenhanced tissues. To control for the staining seen in the unenhanced whole tissue following infusion of 1.9 nm and 15 nm nanogold, 1x PBS was infused into the lateral cerebral ventricle at a rate of 1 μl/min. (A-D) are tissues taken 15 min post 1x PBS infusion. (A) Brain, (B) Subarachnoid space (SAS) of the brain stem, (C) spinal nerve root with the dotted lines marking an example of a nerve root, and (D) trigeminal nerve with dotted lined marking the transition from central to peripheral nervous system. (E-H) are tissues taken 4 hr post 1x PBS infusion. (E) Brain, (F) Subarachnoid space (SAS) of the brain stem, (G) spinal nerve root with dotted lines marking an example of a nerve root, and (H) trigeminal nerve with dotted lines marking the transition from central to peripheral nervous system. 1x PBS tissues are unstained compared to nanoprobe infused animals. N= 3.

**Fig. S5.**
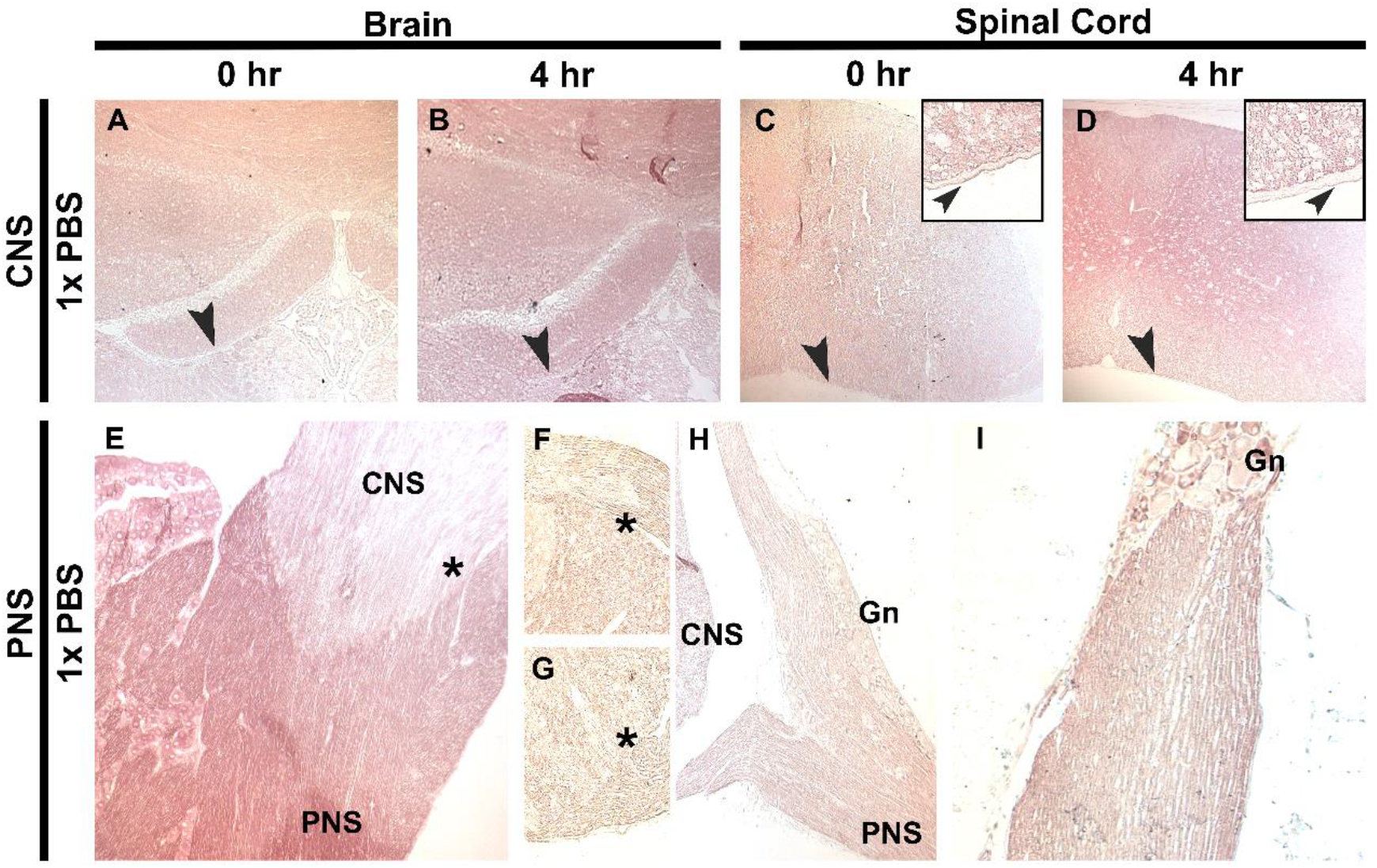
Control 1x PBS histological sections following gold enhance staining. To control for the staining seen in the enhanced histological sections following infusion of 1.9 nm and 15 nm nanogold, 1x PBS was infused into the lateral cerebral ventricle at a rate of 1 μl/min and enhanced for 5 min. (A-D) are sections taken from Central nervous system tissues (CNS). (A) third ventricle and intraventricular foramen 15 min post 1x PBS infusion. (B) third ventricle and intraventricular foramen 4 hr post infusion. Arrowheads mark the unstained intraventricular foramen to compare to that seen in the nanoprobe infused animals. (C) spinal cord 15 mins post 1x PBS infusion, with a high mag (63x) inlay of clear meninges. (D) spinal cord 4 hr post 1x PBS infusion, with a high mag (63x) inlay of clear meninges. Arrowheads denote the spinal meninges. Meninges are clear compared to the 1.9 nm meninges at 15 min and 4 hr post infusion and 15 nm 15 min post infusion. (E-I) are sections taken from the peripheral nervous system (PNS) 4 hr post 1x PBS infusion to correlate with the images in figure 4. “*” marks the transition in all panels. Ganglion are labeled Gn. (E) Trigeminal nerve. (F) dorsal spinal nerve root transition. (G) ventral nerve root transition. (H) dorsal and ventral nerve roots combining with the dorsal root ganglion (Gn) seen. (I) proximal section of the sciatic nerve. (ganglion labeled “Gn). N = 3.

**Fig. S6.**
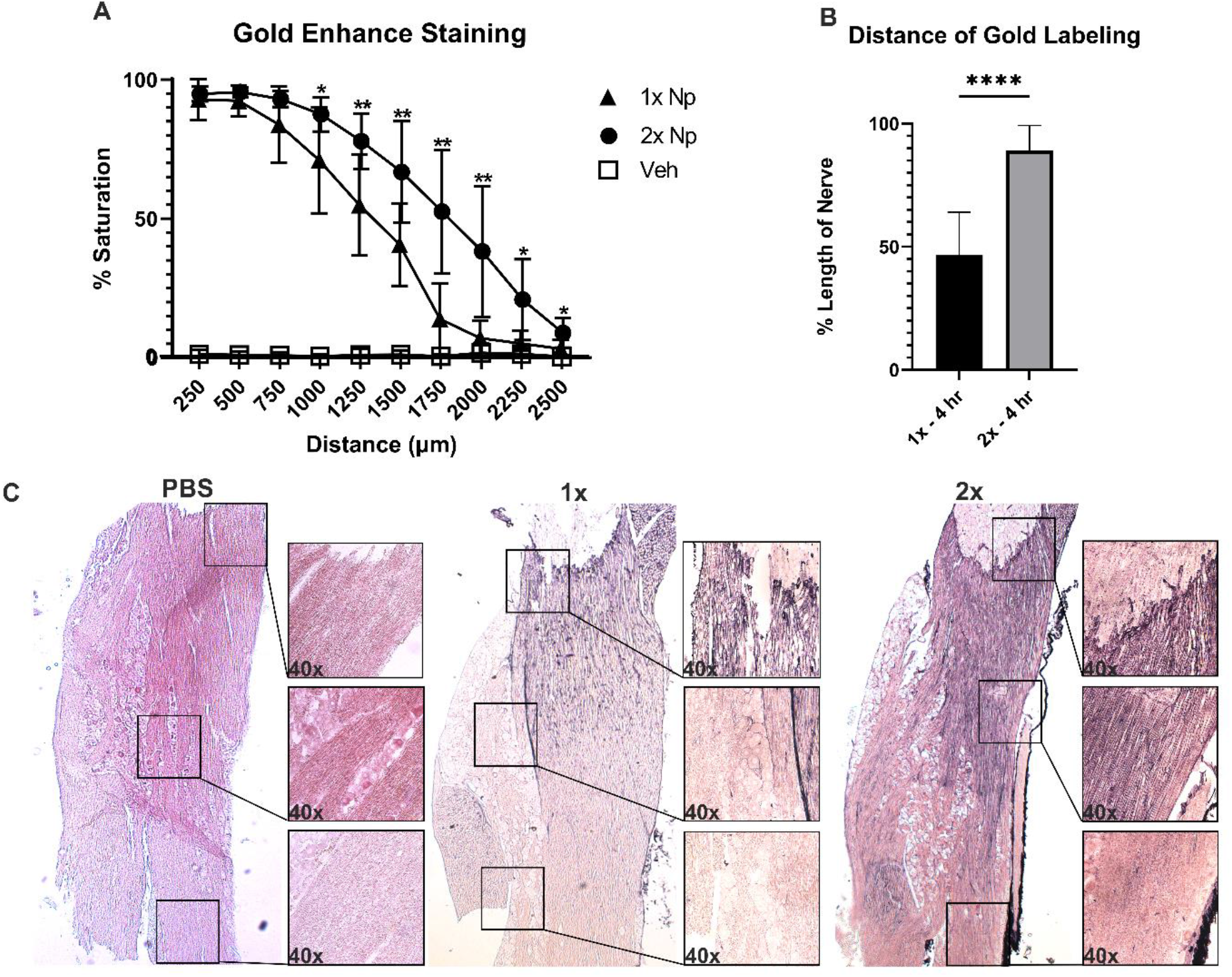
Gold enhanced stain intensity and distance quantification of 1x and 2x concentration. To determine the effect of ICV-infused probe concentration on probe distribution in the PNS, 1 mg (1x) and 2 mg (2x) of 1.9nm nanogold tracer was infused into the lateral cerebral ventricles of mice at a rate of .1 mg/min. Staining intensity of trigeminal nerves was compared. (A) Staining intensity was quantified by measuring pixel density in 1x PBS (Veh), 1x (1x Np) and 2x (2x Np) trigeminal nerves following a 20-minute gold enhancement. Every 250 μm along the nerves were quantified for a total length of 2500 μm. 1x concentration is denoted by a black triangle, 2x concentration by a black circle and 1x PBS by an open square. (B) The distance of gold labeling was measured by the identification of nodal staining in 1x and 2x trigeminal nerves following 20-minute gold enhancement. Doubling the concentration of nanogold doubled the distance of nodal staining seen. (C) 5x image of nerves used in Figure 5 (trigeminal nerves) following 5 minutes of enhancement, with 40x magnification inlays. 1x PBS nerves show no nanogold staining. 1x concentration has a lower level of nanogold staining, for a lesser distance than the 2x concentration. Unpaired t-test was performed to determine statistical significance (* P < 0.05), ** P < 0.01, **** P < 0.0001) Panels have a minimum N=4.

**Fig. S7.**
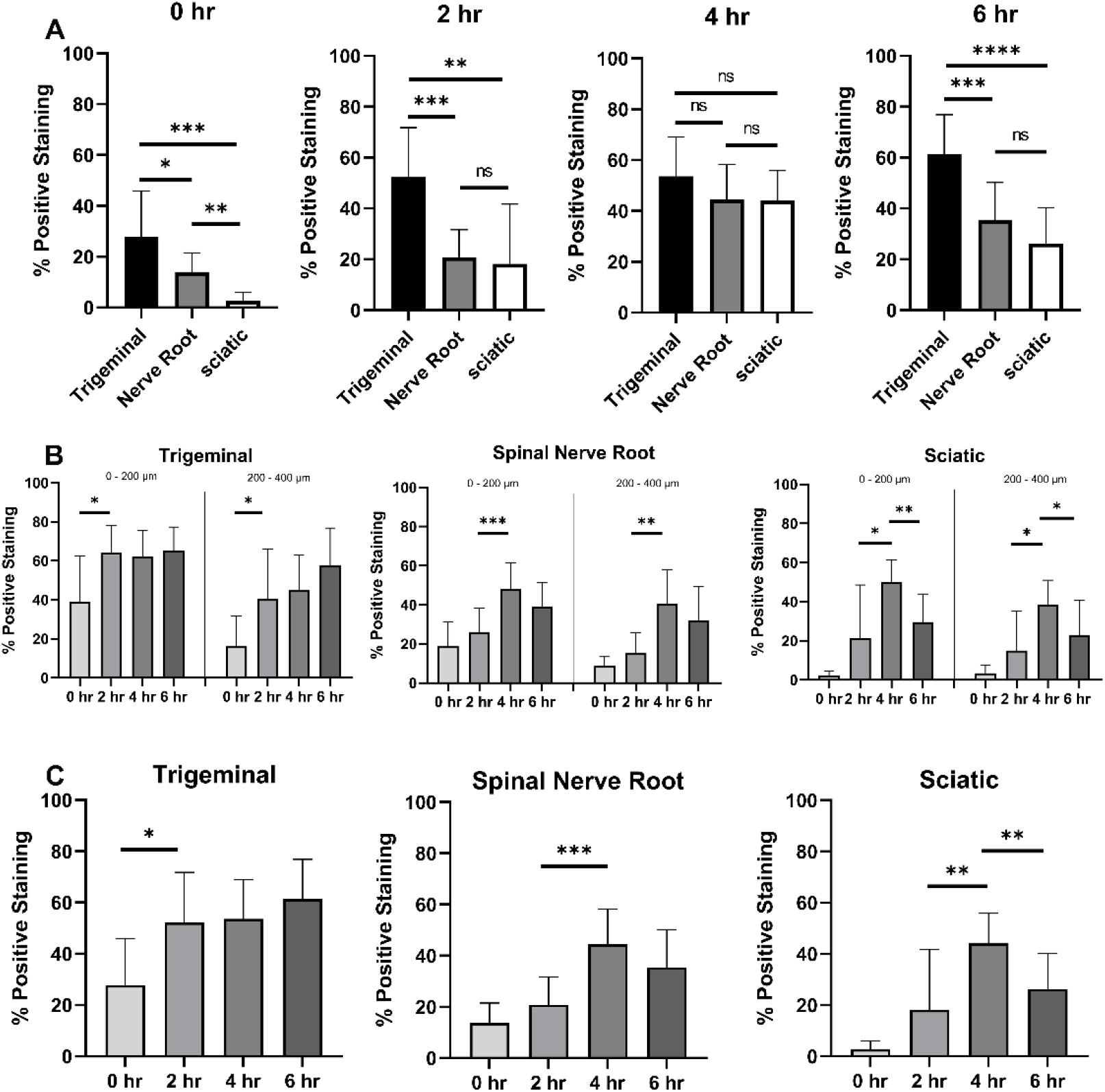
Average nanogold staining of peripheral nervous tissue after 0, 2, 4 and 6 hours. To determine the effect of time on nanoprobe distribution in the peripheral nervous system following intracranial 1.9 nm nanogold tracer infusion, trigeminal nerves, spinal nerve roots, and sciatic nerves were harvested from animals at 15 min (0 hr), 2 hr, 4 hr, and 6 hr after probe infusion (see Figure 3). To quantify analyze probe distribution in these tissue, positive pixel density (positively staining pixels compared to total # pixels in designated area) was quantified using ImageScope. (A) Overall staining at each timepoint was quantified and compared in each tissue. Trigeminal nerve had the highest overall staining seen at each timepoint. (B) Pixel density was measured at 0 – 200μm and 200 – 400μm from the CNS to PNS transition of the trigeminal and spinal nerve root or from the ganglion in the sciatic nerve. At the distance of 0 – 200μm in the trigeminal, maximal staining is reached by 2 hours. From 200 – 400μm, the same level of staining seen in the first 200μm is not reached until 6 hrs post infusion. In the spinal nerve root, and sciatic nerves, the highest level of staining is not seen until 4 hrs post infusion before decreasing again at 6 hrs. (C) The two distances were averaged together at each timepoint. The highest level of staining overall in the trigeminal is at 6 hrs, but 4 hrs in the spinal nerve root and sciatic nerves. Unpaired t-test was performed to determine statistical significance (* P < 0.05, ** P < 0.01, *** P < 0.001).

**Fig. S8.**
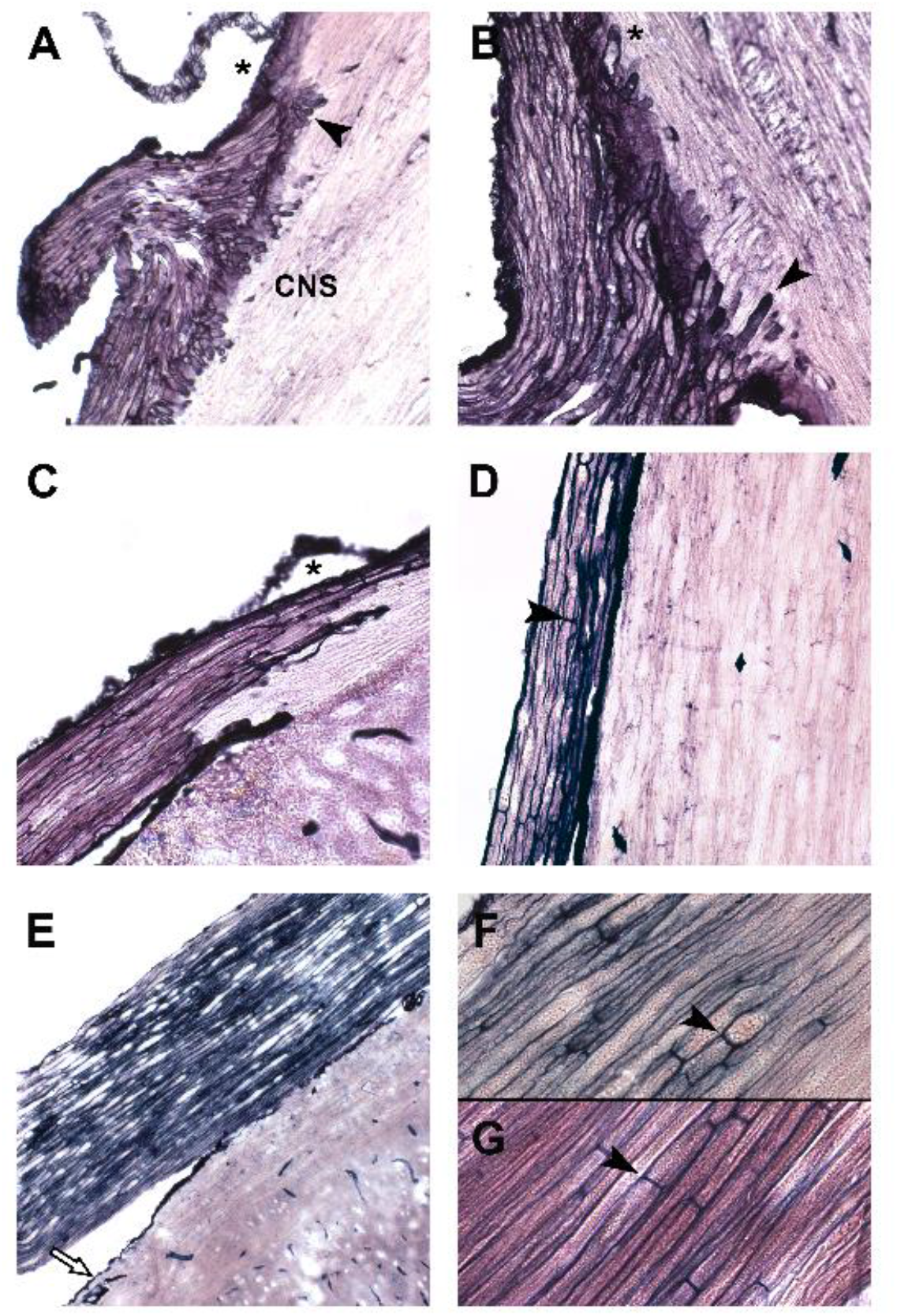
CSF flows through spinal nerves in a time and distance dependent manner. To assess the effect of time on staining in spinal nerves, nanogold was infused into the cerebral lateral ventricle. Spinal cords and nerve roots were collected at 2, 4, or 6 hours (hr) post infusion at cervical, thoracic, or lumbar spinal positions. Tissue sections were GoldEnhanced. (A) Protruding cervical spinal nerve root 2 hr after nanogold infusion; “*” marks branch point for protruding nerve in all panels. (B) High magnification image of cervical nerve root showing nanogold surrounding individual axons (arrowhead). (C) Dorsal thoracic nerve root from nanogold-injected animal 4 hr after ventricular injection. Probe is concentrated in nerve root. (D) Spinal cord/spinal root in 4 hr nanogold animal; nanogold is present within the endoneurium, revealing Nodes of Ranvier (white arrowhead). (E) Spinal cord with branching lumbar nerve root 6 hr after nanogold infusion into ventricle; peripheral neural tissue is labeled substantially more than spinal cord (open arrow marks CNS meninges). (F) Sciatic 6 hr after infusion of 1 mg (1X) nanogold into lateral ventricle; arrowhead marks Nodes of Ranvier in (F) and (G). (G) Sciatic nerve 4 hr after 2X (2 mg) nanogold injection into the lateral ventricle. In high magnification images (F) and (G), gold is detectable in the endoneurium surrounding individual axons and encircling Nodes of Ranvier. Panels have a minimum N=4.

**Fig. S9.**
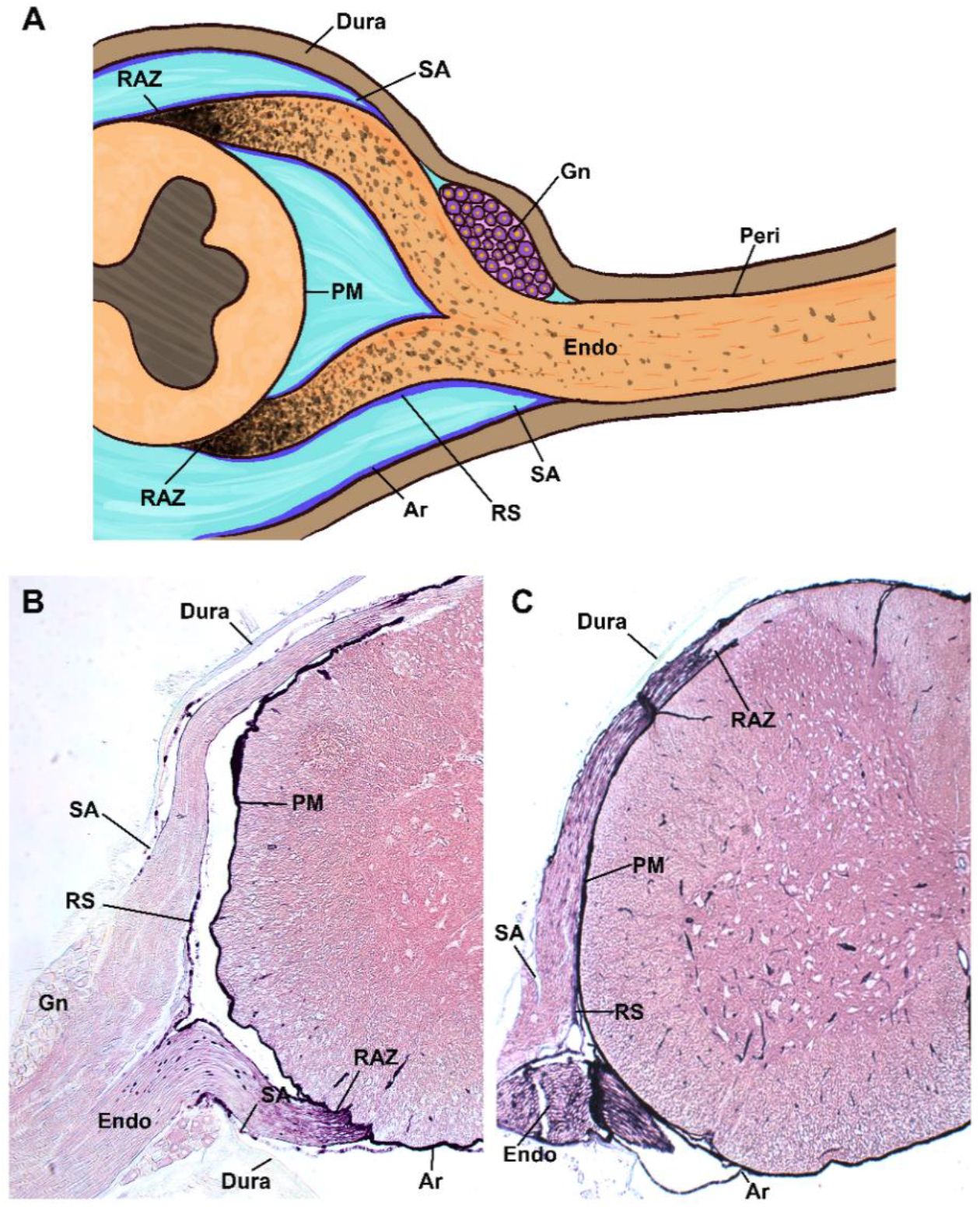
CSF solute enters PNS at nerve root attachment zone. 1.9nm nanogold tracer was infused into the lateral cerebral ventricle of fully anesthetized mice. CNS and PNS tissues were harvested (A) 15 min or (B) 6 hr post infusion and processed with GoldEnhance. (C) Schematic of CNS/PNS spinal junction with structures labeled as: RAZ – root attachment zone; PM – pia mater; Ar – arachnoid mater; DM – dura mater; SA – subarachnoid angle; RS – root sheath; Endo – endoneurium/endoneurial region; Gn – ganglion.

**Fig. S10.**
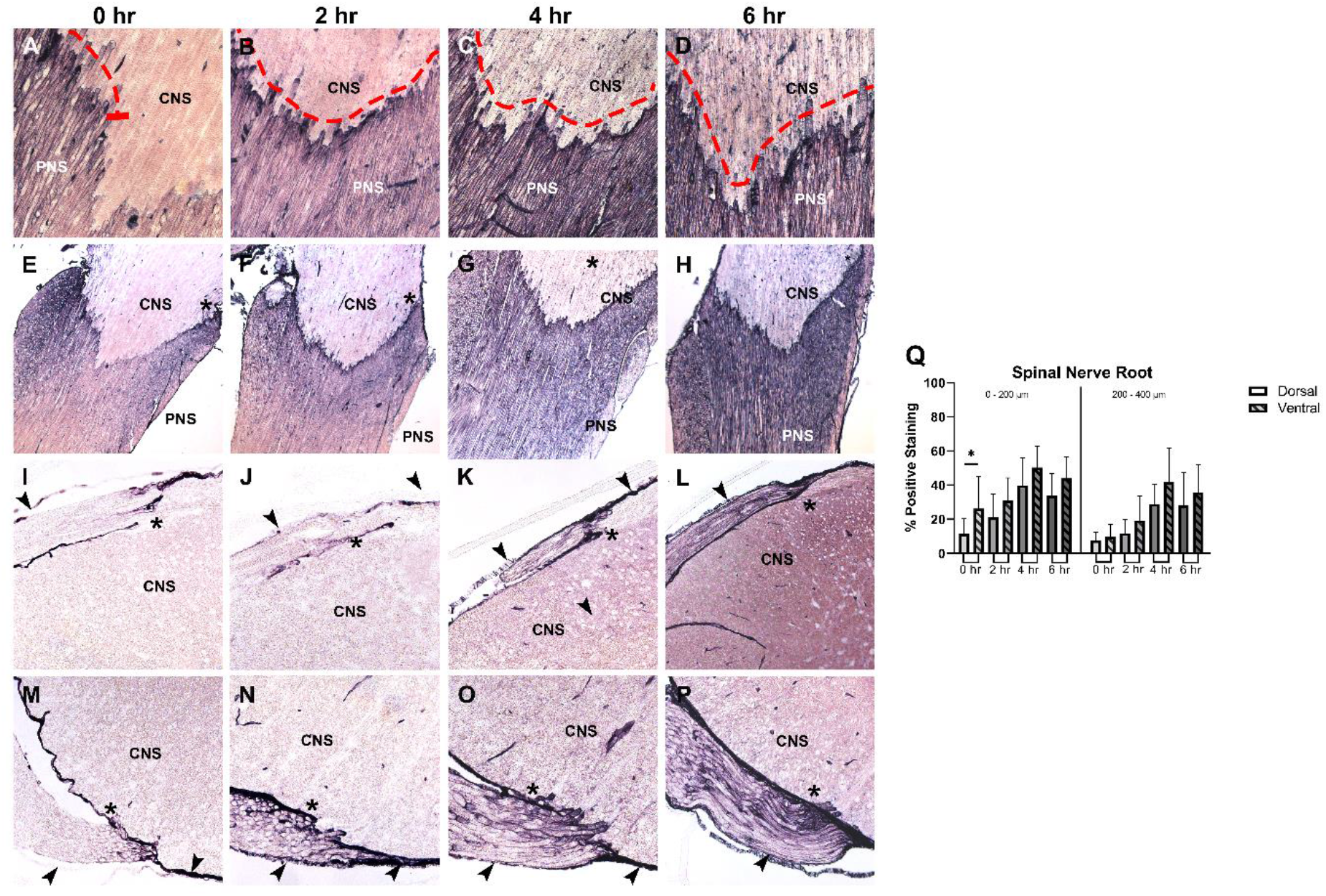
CSF solute accumulates in PNS at CNS/PNS junction. 1.9nm nanogold tracer was infused into the cerebral lateral ventricle. PNS tissues were collected at 15 min, 2, 4, or 6 hours (hr) post infusion, and then cryosections were GoldEnhanced. (A – D) Trigeminal nerves from animals (B) 15 min, (C) 2 hr, (D) 4 hr, or (E) 6 hr post-nanoprobe injection; gold accumulates at CNS-PNS transition as demarcated with red dotted line. (E-H) Low magnification images of trigeminal nerves (E) 15 min, (F) 2 hr, (G) 4 hr, or (H) 6 hr post-nanoprobe injection. CNS to PNS junction is demarcated by “*”. (I-L) Dorsal thoracic nerve root from animals (I) 15 min, (J) 2 hr, (K) 4 hr, or (L) 6 hr post-nanoprobe injection after ventricular injection. Probe is concentrated in nerve roots. Closed arrow marks CNS meninges and PNS connective tissue coverings. (M-P) Ventral spinal nerves roots (M) 15 min, (N) 2 hr, (O) 4 hr, or (P) 6 hr post-nanoprobe injection. Nanogold is present within the endoneurium, revealing Nodes of Ranvier (open arrowheads). (Q) Comparison of dorsal vs. ventral staining was conducted through quantification of positive pixel density (# positive staining pixels per total pixel count in given area) using ImageScope. Unpaired t-test was performed to determine statistical significance (** P < 0.01). Panels have a minimum N=4.

**Fig. S11.**
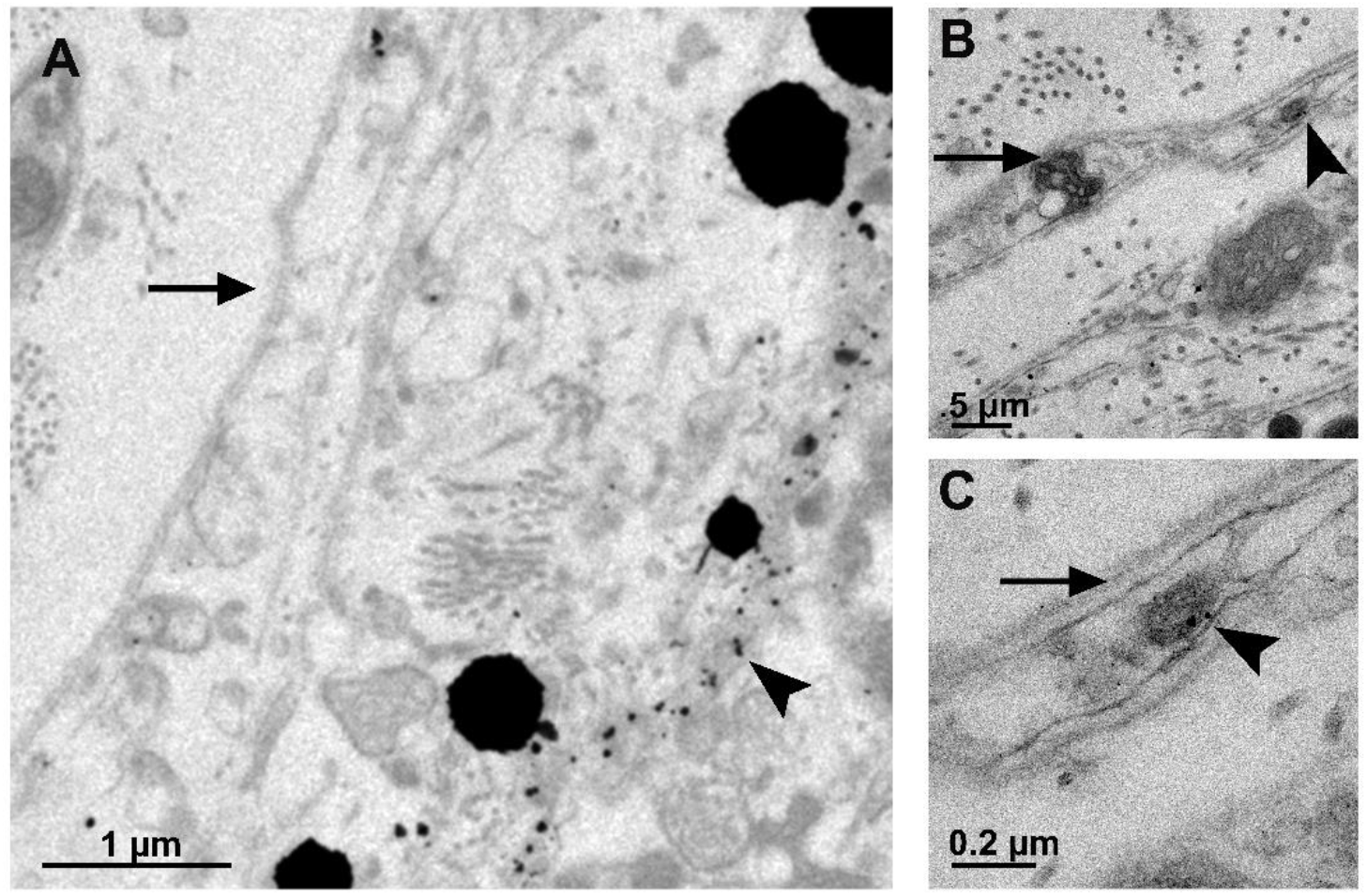
Perineurial cells contain lower levels of nanogold equivalent to CNS levels of staining. All cells of the PNS stain to some degree with nanogold. In order to determine the precise subcellular localization of nanoprobe in 1.9 nm nanogold tracer ICV-infused mice, tissue sections were harvested 4 hr after probe infusion and processed with GoldEnahnce EM for 2 min. Although the perineurium stains the heaviest by light microscopy, electron microscopy reveals minimal nanogold staining within the perineurial cells. (A) A perineurial cell (arrow) identified by the presence of a basement membrane has minimal staining, opposed to the highly stained fibroblast (arrowhead) identified by the lack of a basement membrane. (B) Perineurial cell (arrow) located away from a highly stained fibroblast, with minimal nanogold staining (arrowhead). (C) High magnification of the perineurial cell (arrow), indicating intracellular staining (arrowhead).

**Fig. S12.**
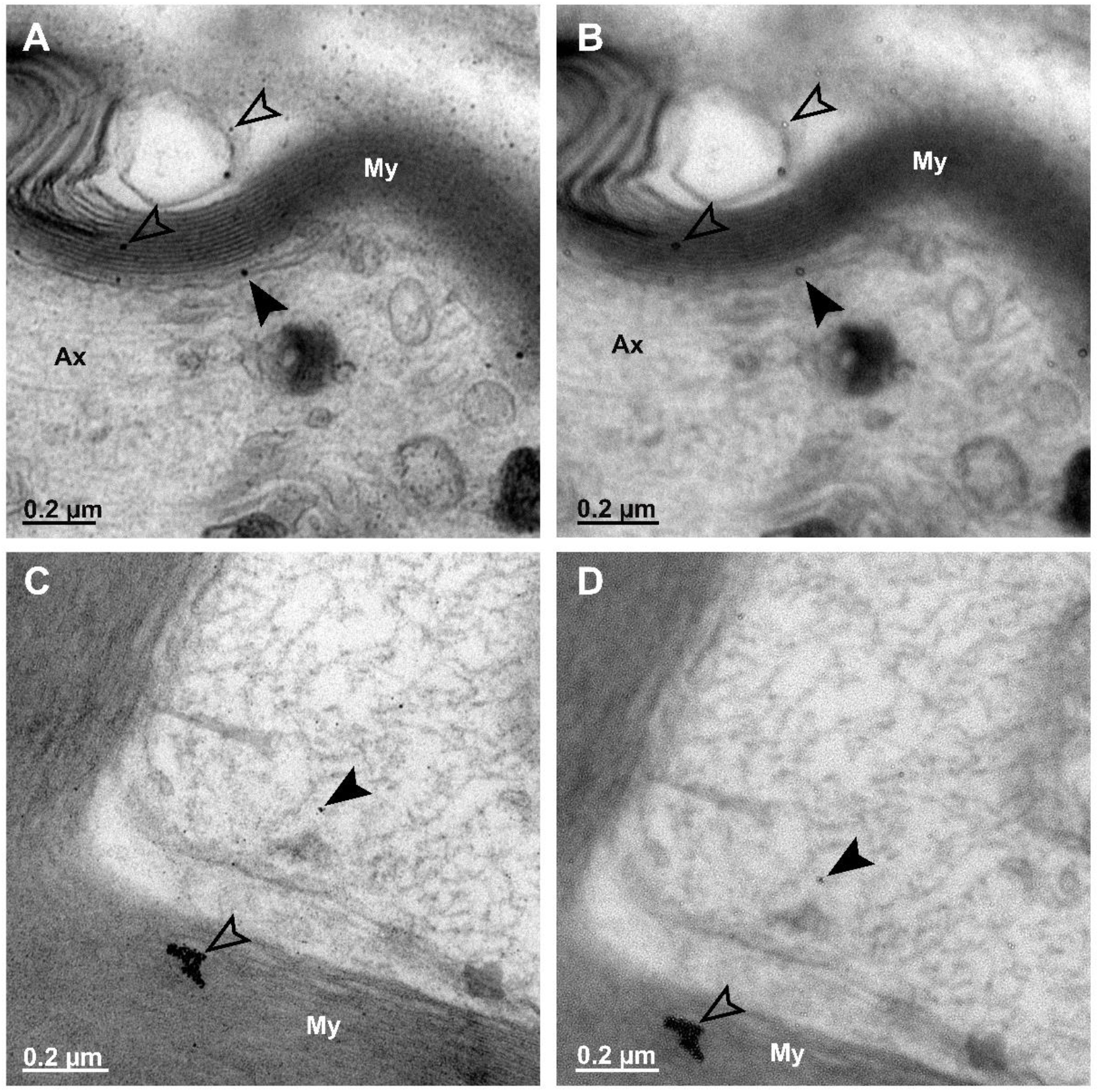
Electron reflection verifies nanogold by electron microscopy. Electron microscopy (EM) was used to determine the cellular localization of nanoprobe in central and peripheral nervous system tissues from mice with 1.9 nm nanogold tracer infused into the lateral cerebral ventricle; tissue sections were harvested 4 hr after probe infusion and processed with GoldEnahnce EM for 2 min. Electron reflection was used to distinguish gold aggregates from endogenous structures. Lumbar nerve root (A,B) and sciatic nerve (C,D) 4 hours after 2 mg/10 μl ventricular infusion of nanogold and 5 min enhancement with GoldEnhance EM. Electrons reflect off the gold nanoparticle producing a halo, which allows for the verification of the presence of nanogold. (A) Schwann cell and axon (Ax) from a lumbar spinal nerve root with the presence of nanogold within the Schwann cell, myelin (My) (open arrowheads), and in the axon (arrowheads). (B) The halos produced by refraction off the same nanogold particles as (A) were identified. (C) A clump of nanogold located in the myelin (My) of sciatic nerve (open arrowhead), and within the axon that the myelin surrounds (arrowhead). (D) Reflection of the clump of nanogold and an example of axonal nanogold, verifying the identity as nanogold.

**Fig. S13.**
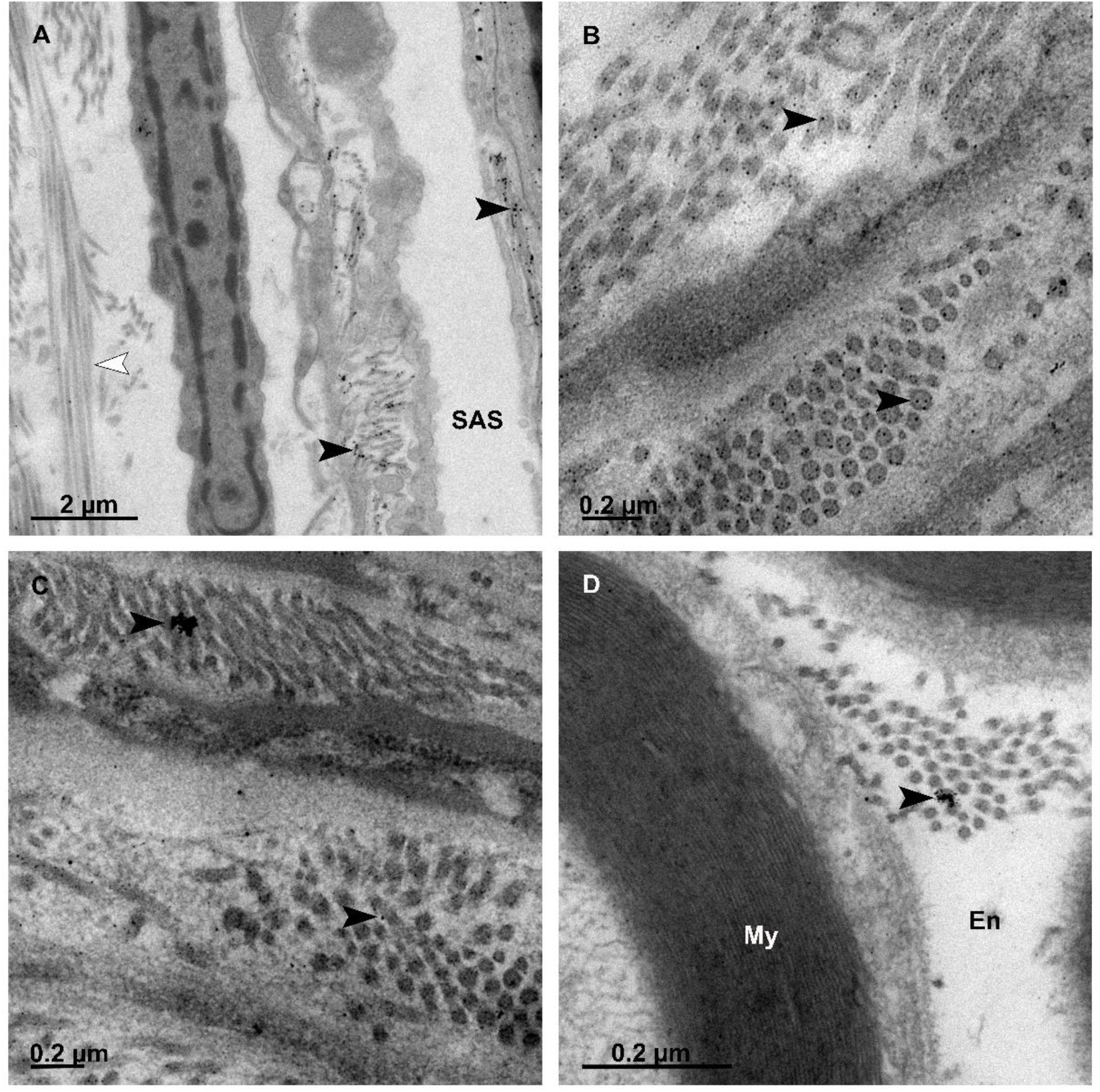
Nanogold accumulates on collagen fibers in the peripheral nervous system (PNS). Free nanoprobe tends to be washed out through normal tissue processing. The nanogold visualized is a result of the particles binding to intracellular and extracellular structures. The bulk of peripheral CSF flow appears to be extracellular identified by an increase in nanogold on collagen fibers that make up the extracellular spaces of peripheral nerves. Electron microscopy identifies the presence of nanogold on collagen fibers. (A) The collagen located between cell layers in the arachnoid and pia meninges (black arrowhead) of a spinal cord have nanogold present, opposed to the subarachnoid space (SAS) and collagen located in the dura (white arrowhead), which is free of nanogold. (B, C) Collagen located between perineurial cell layers in a sciatic nerve, and within the endoneurium (D) are positive for nanogold (black arrowhead).

**Fig. S14.**
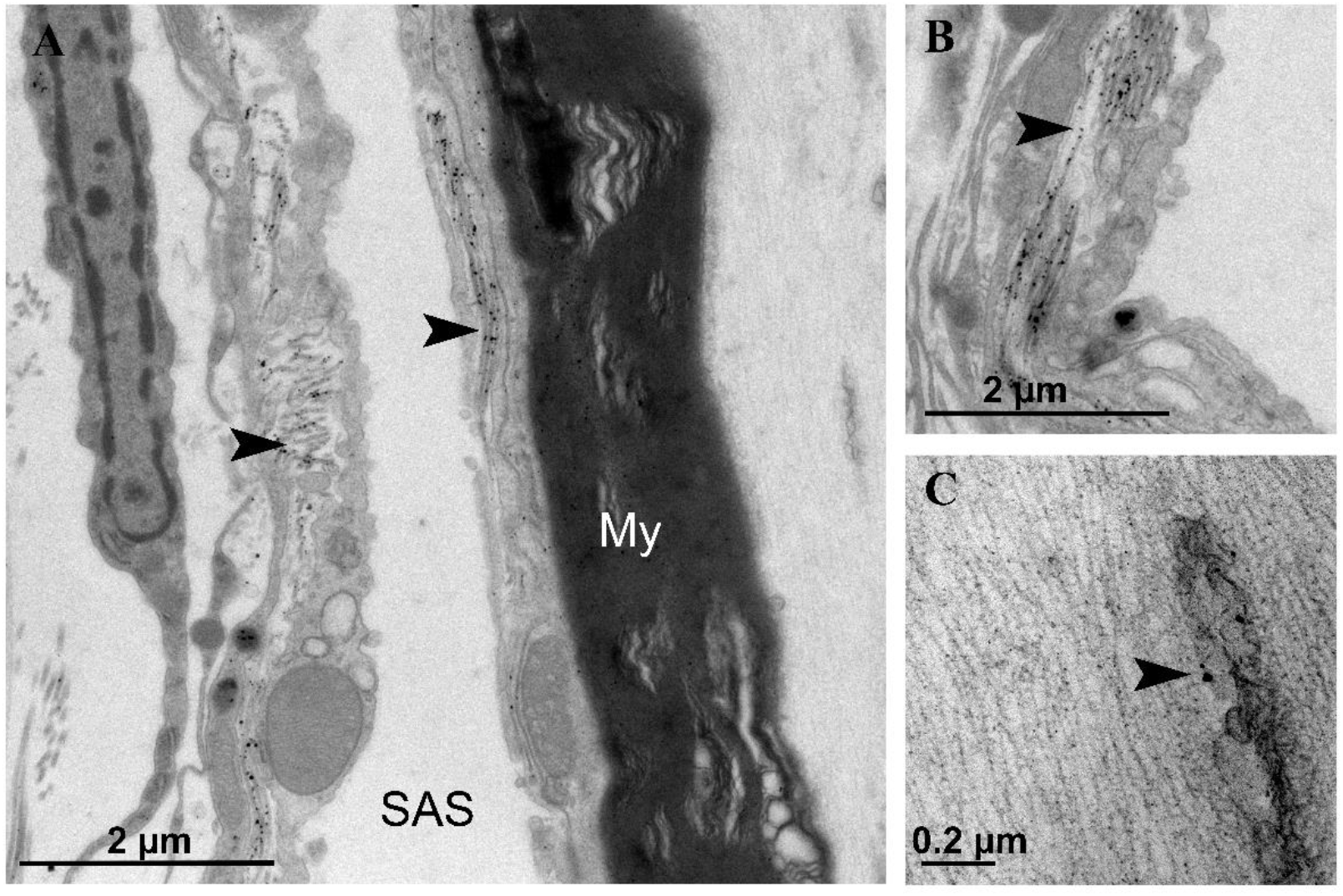
Electron microscopy of central nervous system tissue. Nerves fixed with Trumps fixative, stained with gold enhancement EM for 5 minutes. Electron microscopy of spinal cord 4 hours after nanogold infusion, 2 mg/10 μl infusion of nanogold into the lateral ventricle. (A) Meninges of spinal cord with nanogold amongst the collagen between arachnoid and pia cell layers (arrowhead). The subarachnoid space (known route of CSF flow) does not have nanogold deposition likely due to wash out during processing. The gold nanoparticle is only seen bonded to a structure (collagen in the endoneurium). Due to the absence of extracellular matrix (ECM) in the subarachnoid space, the nanogold is unable to be fixed during tissue processing. Nanogold is seen between the layers of the meninges in the ECM. (B) Magnified view of nanogold between cells in the arachnoid meninges (arrowhead). (C) Similar to what is seen in the PNS, nanogold is located amongst neurofilament in axon found within the central nervous system, albeit at a lower concentration.

**Fig. S15.**
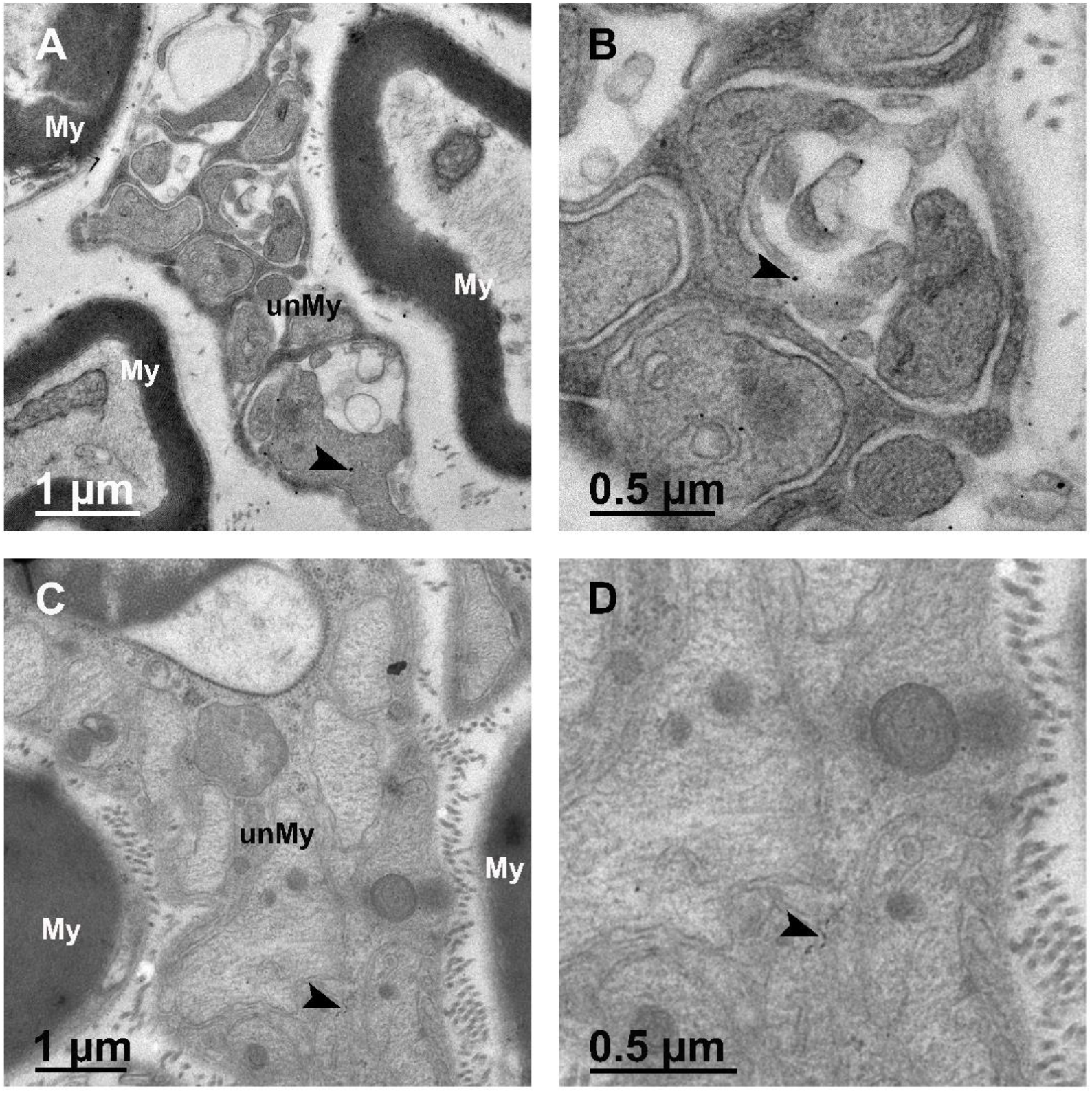
Unmyelinated Schwann cells exhibit lower levels of intracellular nanogold. To assess the distribution of nanogold within peripheral nerves, myelinated and unmyelinated Schwann cells/axons were compared. Nerves were harvested 4 hr post ICV infusion of 2mg/10μl 1.9nm nanogold. Sections were processed with gold enhancement for 2 minutes and then imaged. A, B are examples from a trigeminal nerve, B is a high magnification image of A. C, D are examples from sciatic nerve, D is a high magnification image of C. Less nanogold was identified in unmyelinated Schwann cells and axons than in myelinated (Fig. 10). Arrowheads mark examples of nanogold in all panels.

**Fig. S16.**
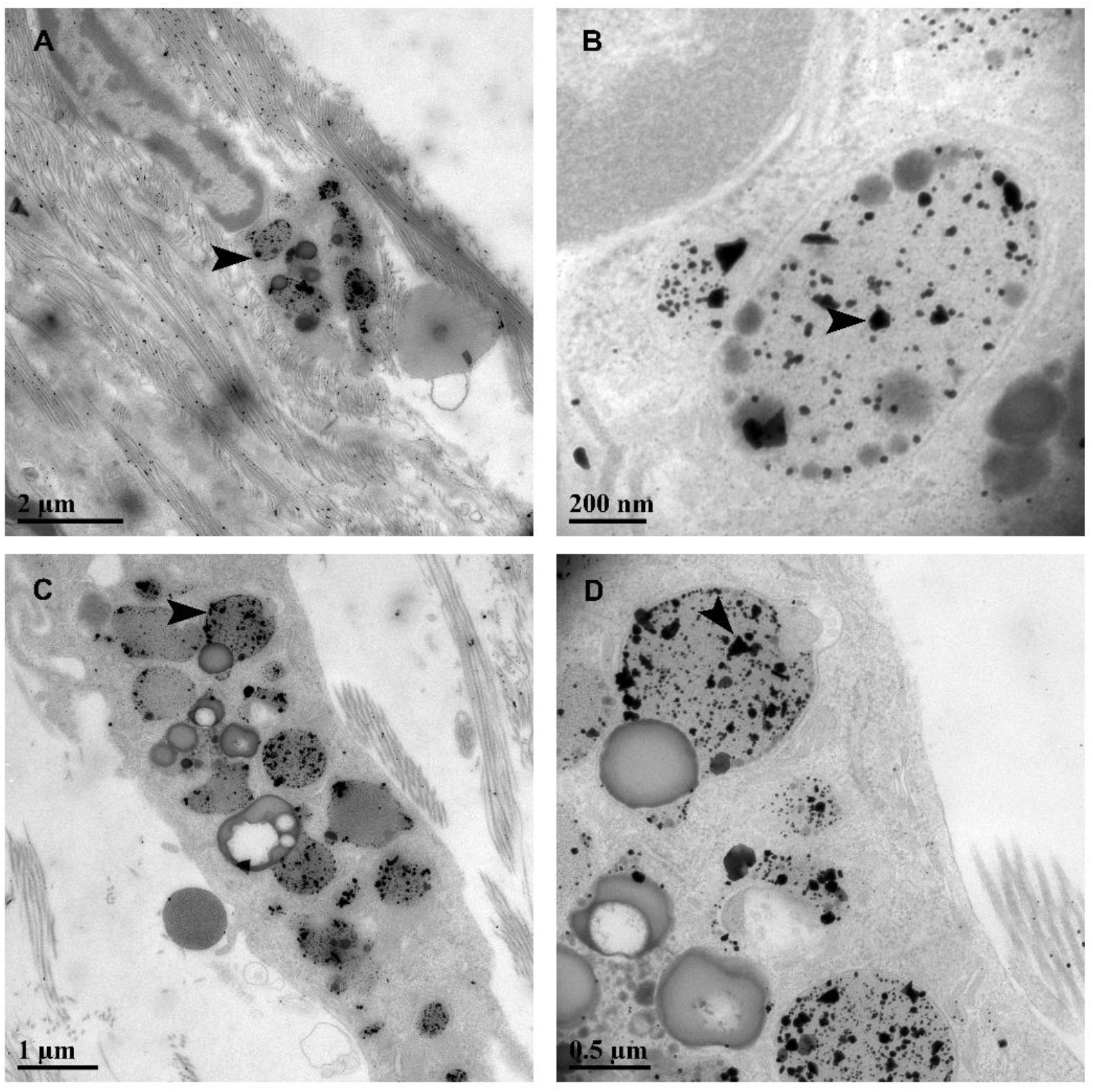
Nanogold is phagocytosed in peripheral nerves. To determine the subcellular localization patterns in the PNS following ICV infusion of 1.9nm nanogold tracer, spinal nerve roots were harvested 4 hr after nanoprobe infusion. Sections were processed with GoldEnhance EM for 2 min and then imaged. Peripheral nerve macrophages were seen in the peri-and endoneurium with nanogold located intracellularly. (A, B) are located within the perineurium of a spinal nerve root, with arrowheads marking nanogold located in vesicles of (A) low magnification and (B) high magnification. (C,D) peripheral nerve macrophage located within the endoneurium of a spinal nerve root. Arrowheads mark nanogold located in vesicles of (C) low magnification and (D) high magnification.

**Fig. S17.**
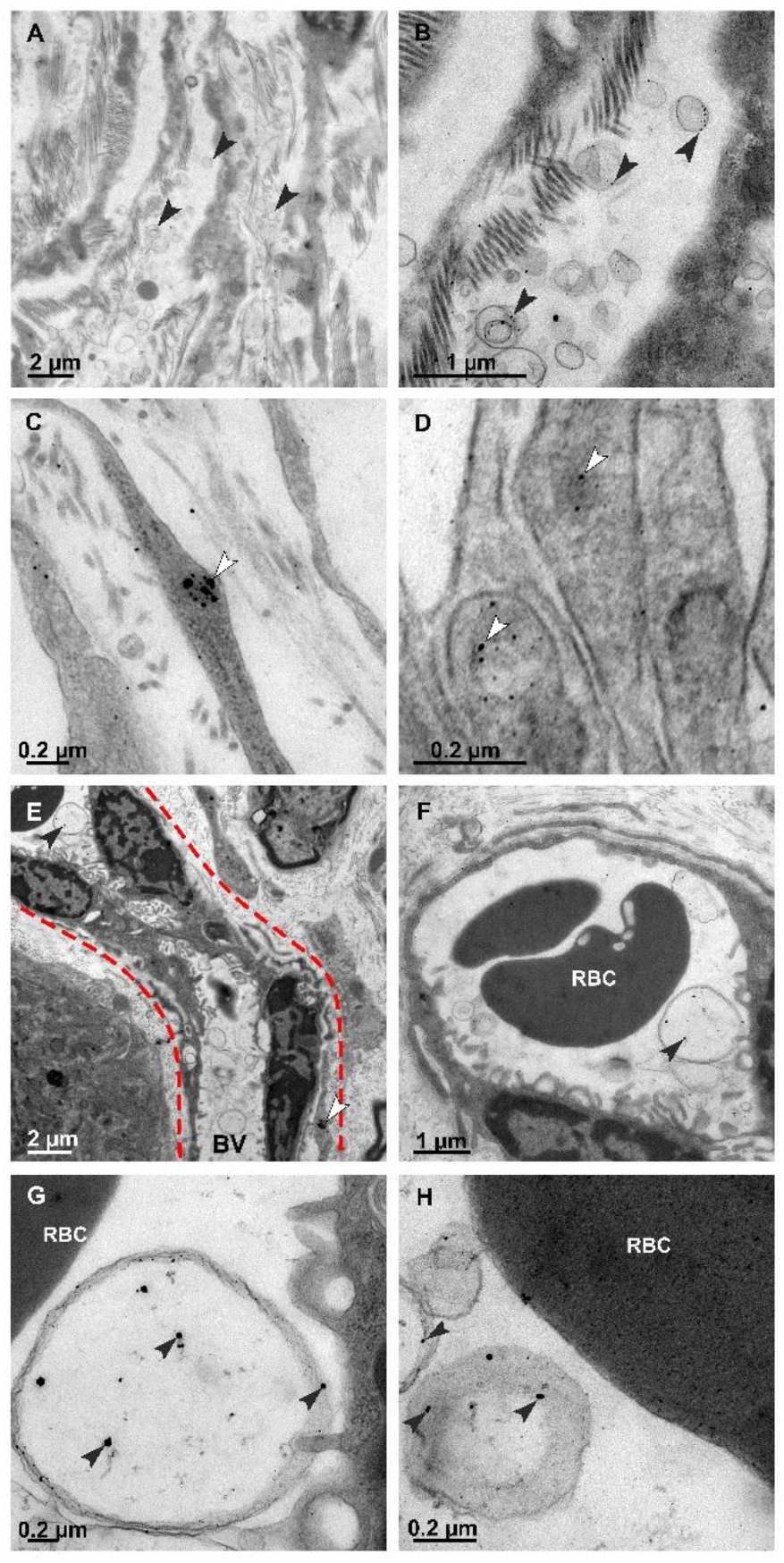
Nanogold accumulates in extracellular and intracellular vesicles. To determine the location of nanoprobe accumulation/subcellular location of nanoprobe distribution following ICV injection, 1.9 nm nanogold tracer was via ICV into fully anesthetized C57BL/6 mice. Tissues were harvested at 4 hrs post infusion and were developed with GoldEnhance EM for 2 min. (A-D) Extracellular vesicles (black arrowheads) and intracellular vesicles (white arrowheads) within the perineurium of spinal nerve roots. (A) Many vesicles are seen between cell layers within the perineurium, often containing nanogold (B). (C,D) Higher magnification reveals nanogold accumulating in vesicular structures within cells of the perineurium (white arrowhead) and between cells (black arrowhead). (E-H) Vesicles located extracellularly (black arrowhead) and intracellular (white arrowhead) in blood vessels located in the spinal nerve roots. (E,F) Low magnification image of a blood vessel within the endoneurium, with vesicles located in the lumen (black arrowhead) and within the endothelial cells (white arrowhead). (G,H) High magnification image of (F) shows nanogold located within the vesicles of the blood vessel, directly adjacent to a red blood cell (RBC).

**Fig. S18.**
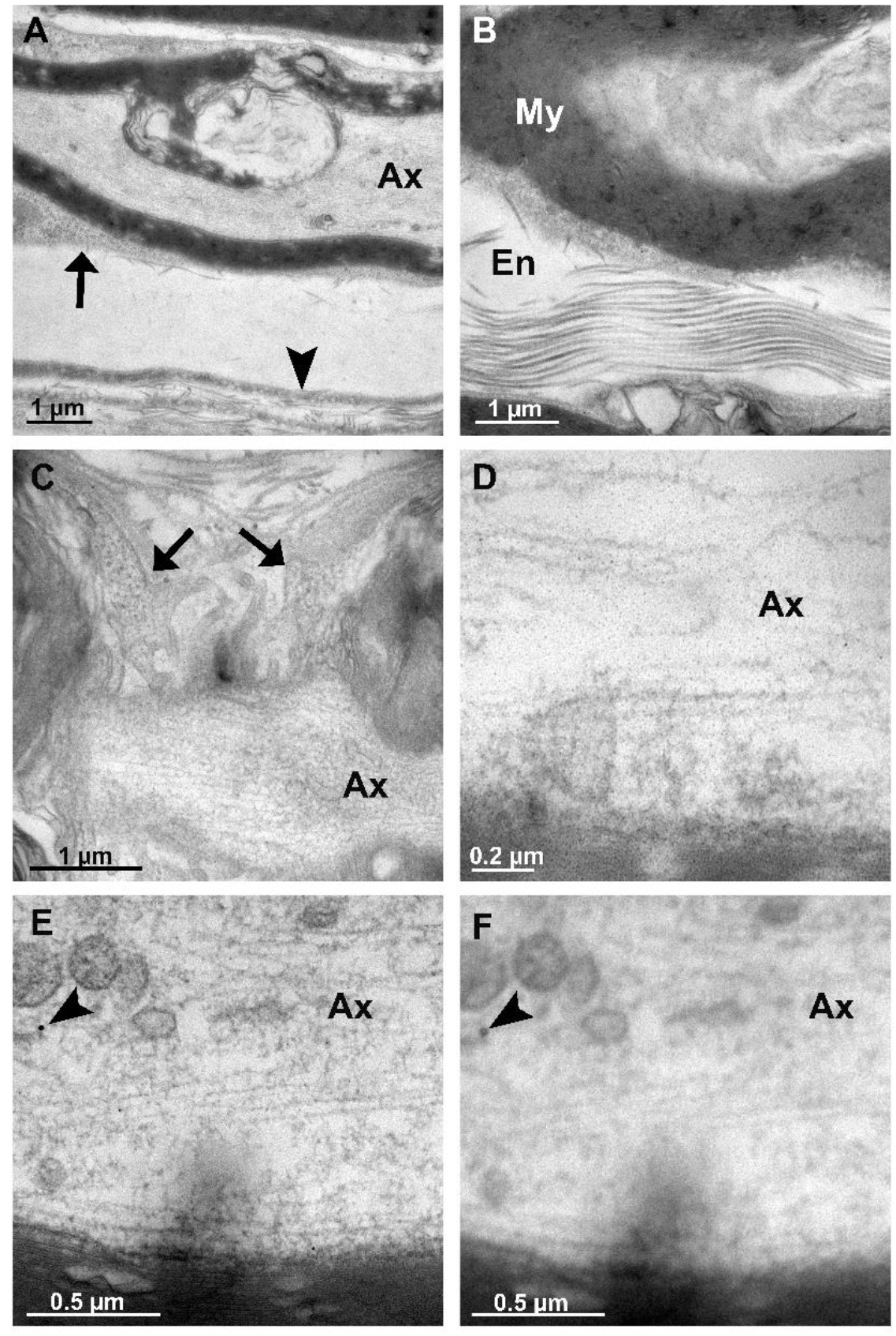
Electron microscopy of trigeminal nerves from control animals. To control for background staining, electron microscopy (EM) of uninjected trigeminal control nerve were imaged following 5 minutes processing with GoldEnhance. The uninjected nerve lacks nanogold labeling in the perineurium, endoneurium, Schwann cell, and axon. (A) No nanogold is seen in the perineurium (arrowhead) or in the Schwann cell/axon (arrow) in an uninjected trigeminal nerve, contrasting the high staining seen in Figure 4b, h. (B) EM identifies a lack of staining in the endoneurium (En). In contrast to Figure 4e-g, collagen in the endoneurium is negative for nanogold. (C) Ultrastructure of a Node of Ranvier is negative for nanogold in both the Schwann cell cytoplasm (arrows) and in the axon (Ax). This is contrasting the high degree of nanogold staining seen by light microscopy in Figure 1c. (D) Neurofilament within the axon (Ax) of an uninjected trigeminal nerve is negative for nanogold staining (compare to Figure 4h-j). (E) Maximal level of background staining seen in an uninjected axon showing a small gold particle, verified by refraction (F). Refraction causes a halo to be produced around the gold particle, allowing verification of gold in tissue in contrast to endogenous proteins that do not refract light.

**Fig. S19.**
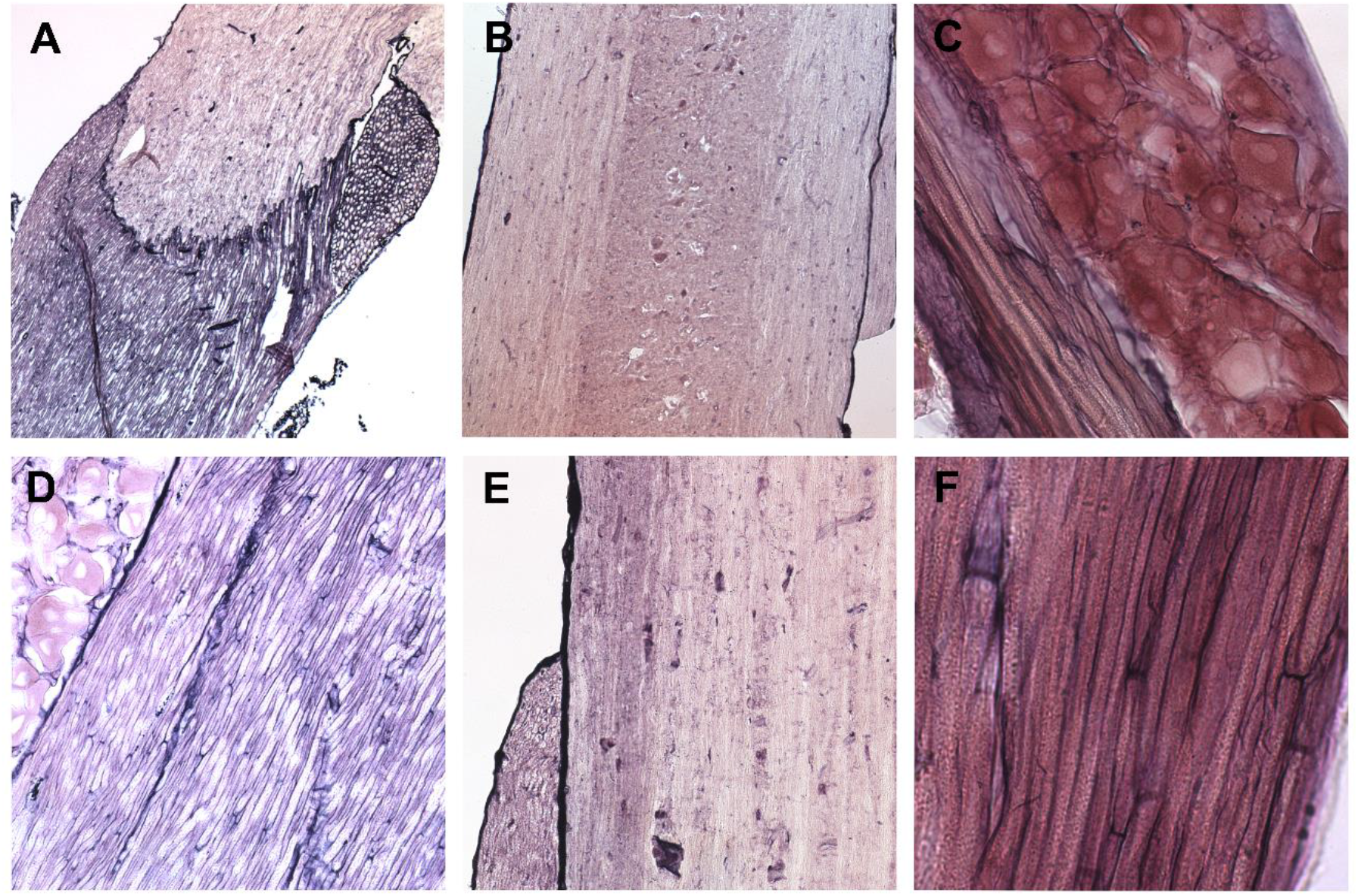
Light Microscopy of the nerves used for electron microscopy. Nerves opposite of those used for electron microscopy were fixed with Trumps fixative, stained with gold enhancement LM for 5 minutes. (A, D) Trigeminal nerve, (B, E) spinal cord, (C, F) lumbar dorsal root ganglion and sciatic nerve taken from mice used for electron microscopy at 4hr post infusion of 2 mg/10 μl 1.9nm nanogold tracer into the lateral ventricle. (A) Trigeminal nerve indicates high concentration of nanogold on the PNS side of the transition zone (*), while minimal intensity seen on the CNS side. Continuous staining seen from the CNS meninges to the Peripheral nerve layers (arrow). (D) Further from the transition zone of the trigeminal nerve, nano gold continues along the perineurium (arrow), endoneurium and accumulating at the nodes of Ranvier (arrowhead). (B) meninges of spinal cord are highly labeled (open arrow), while the parenchyma of the spinal cord has minimal staining (open arrowhead). (E) Higher magnification of spinal cord with labeled spinal meninges. (C, F) Endoneurial staining seen with high nanogold deposition at the nodes of Ranvier (arrowheads).

